# Comparative Genomics of Bovine and Human *Fusobacterium necrophorum* Strains Reveal Subspecies- and Host-associated Differences in Virulence and Antimicrobial Resistance

**DOI:** 10.64898/2026.04.22.720273

**Authors:** Justine Kilama, Devin B. Holman, Mina Abbasi, Raghavendra G. Amachawadi, T.G. Nagaraja, Carl R. Dahlen, Samat Amat

## Abstract

*Fusobacterium necrophorum* (FN) is an important opportunistic pathogen implicated in necrotizing infections, including liver abscesses, calf diphtheria, metritis, and foot rot in cattle, and tonsillopharyngitis in humans. However, FN also exists as a commensal member of the bovine reproductive microbiota with potential negative, as in metritis, and even positive associations with pregnancy outcomes. The genomic features that enable FN to colonize diverse hosts and anatomical niches as either a commensal or a pathogen is poorly understood. We addressed these knowledge gaps by performing comparative genomic analysis of 137 FN strains (80 newly sequenced, 57 publicly available) from clinical and non-clinical sources across human and bovine hosts. We investigated the pangenome structure, virulence gene repertoire, antimicrobial resistance genes (ARG) prevalence, as well as host-and subspecies-associated genomic signatures of two FN subspecies: subsp. *necrophorum* (FNN) and subsp. *funduliforme* (FNF). Comparative genomics revealed an open pangenome with high accessory diversity, and phylogenetic analysis separated the strains into two distinct subspecies clades. Functional profiling revealed substantial metabolic divergence between subspecies, with FNN showing higher prevalence of carbohydrate transport systems and advanced glycation-related pathways, while FNF showed enrichment in threonate metabolism and hemolysin-related systems. Virulence gene analysis identified 84 variants across multiple functional categories with subspecies- and host-specific distributions. Antimicrobial resistance genes, primarily tetracycline resistance genes [*tet*(O), *tet*(M), *tet*(40)] and the macrolide resistance gene *erm*(B), were detected in 22.6% of strains, with higher prevalence in bovine than human strains. Overall, our results suggest that pathogenic potential of FN appears to be determined by the interplay between an open pangenome, subspecies-specific metabolic and virulence repertoires, host-associated adaptation, and niche specialization.

**IMPORTANCE:** *Fusobacterium necrophorum,* comprising two subspecies, *necrophorum* (FNN) and *funduliforme* (FNF), is a major pathogen in cattle and humans, yet it also occurs as a commensal inhabitant in healthy cattle, particularly in the rumen, hindgut, semen, and the female reproductive tract. However, emerging evidence indicates it also occurs as a non-clinical inhabitant, particularly in the bovine reproductive tract, where it may be associated with improved pregnancy outcomes. We conducted a comparative genomic analysis of 137 *F. necrophorum* strains, including FNN (n = 12) and FNF (n = 125), sourced from humans (n=53) and cattle (n=84) across seven anatomical niches spanning both healthy and diseased sources. We identified subspecies- and host-specific metabolic pathways, antimicrobial resistance profiles, and distinct virulence gene distributions that underpin the ecological versatility of *Fusobacterium necrophorum*. Overall, these findings provide a genomic framework for understanding its host adaptation and niche specialization across bovine and human hosts.

## INTRODUCTION

*Fusobacterium necrophorum* is a Gram-negative, pleomorphic, non-motile, non-spore-forming, and aerotolerant anaerobic bacterium traditionally recognized as one of the most common opportunistic pathogens in veterinary medicine, particularly in cattle (Langworth, 1977; Nagaraja and Chengappa, 1998; Nagaraja et al., 2005). *F. necrophorum* is taxonomically divided into two subspecies: subsp. *necrophorum* (FNN) and subsp. *funduliforme* (FNF) (Tadepalli et al., 2009). *F. necrophorum* causes a wide spectrum of necrotizing infections across multiple livestock and wildlife species, as well as in humans (Nagaraja et al., 2005). In cattle, it contributes to economically important diseases such as liver abscesses, calf diphtheria, metritis, and foot rot (Tadepalli et al., 2009; Amachawadi and Nagaraja, 2022), whereas in humans it is associated with tonsillopharyngitis, including tonsillitis and tonsillar abscesses and in rare cases, Lemierre’s syndrome (Riordan, 2007; Dalen and Mekhail, 2015). In dairy cows, the clinical manifestations lead to substantial economic losses and welfare concerns in North American cattle production (Valerio et al., 2020; Taylor et al., 2025).

However, emerging evidence from our recent studies challenges the conventional view of *F. necrophorum* as mainly pathogenic bacterial agent in reproductive tract (Webb et al., 2023b; Kilama et al., 2025). In a 16S rRNA sequencing study, we observed a positive association between *F. necrophorum* abundance and pregnancy success in beef cattle, with cows that became pregnant exhibiting a significantly greater abundance of *F. necrophorum* in the uterine microbiota at artificial insemination compared to those that did not become pregnant (0.63% vs. 0.005%) (Webb et al., 2023b). Similarly, characterization of the seminal microbiota in healthy beef bulls revealed that *Fusobacterium* spp. comprised approximately 26% of the total bacterial community, indicating their prominence within the normal reproductive microbiota (Webb et al., 2023a). This is further supported by targeted culturing and quantitative polymerase chain reactions (qPCR) assays that we conducted previously, which confirmed the presence of viable *F. necrophorum* subspecies in 66.7% of healthy bull seminal samples. These findings indicate that *Fusobacterium* spp. are established as commensals rather than transient contaminants (Kilama et al., 2025). Notably, strains were recovered from multiple reproductive tract niches, including vaginal and uterine samples, consistent with stable colonization of healthy hosts (Kilama et al., 2025). Beyond the reproductive system, *F. necrophorum* is capable of adapting to various anatomical niches, including respiratory, gastrointestinal, and reproductive systems across multiple host species (Nagaraja et al., 2005; Kumar et al., 2013a; Pillai et al., 2019; Pillai et al., 2021; Kilama et al., 2025). In addition, qPCR analyses detected *F*. *necrophorum* in tissues from 260-day-old calf fetus, including placental caruncles, fetal fluids, and meconium (Kilama et al., 2025), suggesting the calf fetus may get exposed to *F*. *necrophorum* prenatally. Taken together, *F. necrophorum* may function as a context-dependent member of the microbiome, challenging its traditional classification as a mainly pathogenic organism.

Despite its wide distribution across diverse anatomical niches within a host and across different hosts, the genomic features that enable *F. necrophorum* to colonize both bovine and human hosts, functioning as either a commensal or a pathogen, remain poorly understood. The two subspecies of *F. necrophorum*, FNN and FNF, are traditionally differentiated through subspecies-specific colony morphology on blood agar, microscopic morphology, and biochemical characteristics (Tan et al., 1994; Nagaraja et al., 2005; Jensen et al., 2008). Definitive molecular identification is achieved using qPCR targeting distinct leukotoxin operon promoter regions, specifically *lktA-n* for FNN and *lktA-f* for FNF (Deters et al., 2024; Kilama et al., 2025). The subspecies FNN is notably more virulent than FNF, owing to its ability to produce higher levels of leukotoxin, and consequently has been isolated more frequently from infections (Pillai et al., 2019; Amachawadi and Nagaraja, 2022; Bista et al., 2022).

Most studies have focused on *F. necrophorum* strains recovered from infected tissues and virulence-associated genes, with relatively little attention given to broader genomic variation between subspecies or to strains colonizing commensal niches (Bicalho et al., 2012; Bista et al., 2022; Wynn et al., 2022; Carrara et al., 2024). Therefore, several key knowledge gaps exist, including: (1) subspecies- and host-associated genomic signatures; (2) genomic features distinguishing clinical from non-clinical strains; (3) patterns of virulence factor distribution across commensal and pathogenic populations, and (4) the occurrence and diversity of antimicrobial resistance genes (ARGs) within the *F. necrophorum*. We hypothesized that subspecies identity, host species, and anatomical niche would be associated with distinct genomic signatures linked to commensal and pathogenic lifestyles. The objectives of the present study were to: (i) characterize the pangenome structure and subspecies-specific genomic differences between FNN and FNF; (ii) identify virulence factor-encoding genes, functional gene content, and genomic adaptations associated with host species and anatomical niche; and (iii) assess the prevalence and distribution of ARGs across ecological niches. To address these objectives, we performed whole-genome sequencing and comparative genomic analyses on 137 *F. necrophorum* genomes from human and bovine hosts, representing both subspecies (FNN and FNF) and spanning seven anatomical niches, including semen, liver (healthy and abscessed), lungs, ruminal fluid, larynx (calf diphtheria), hoof (foot rot), and mediastinal lymph nodes.

## MATERIALS AND METHODS

### Isolate collection and culturing conditions

A total of 80 *Fusobacterium necrophorum* (FN) strains were subjected to whole-genome sequencing in this study, comprising 69 subsp. *funduliforme* (FNF) and 11 subsp. *necrophorum* (FNN). Samples collected by our laboratory (Department of Microbiological Sciences, North Dakota State University) from cattle in North Dakota and from cattle raised by the U.S. Meat Animal Research Center (Clay Center, Nebraska) were submitted to the Department of Diagnostic Medicine and Pathobiology, College of Veterinary Medicine at Kansas State University College of Veterinary Medicine, where strains were cultured and recovered. Additional strains were obtained from the laboratory collection of T. G. Nagaraja, as well as from Raghavendra (Department of Clinical Sciences, College of Veterinary Medicine, Kansas State University) and the North Dakota State University Veterinary Diagnostic Laboratory. Further details on isolate numbers and origins are provided in Table 1. The FNF and FNN strains were isolated from multiple anatomical sites of cattle, include the liver (healthy and abscessed), semen, ruminal fluid, larynx (calf diphtheria), mediastinal lymph nodes, hoof (foot rot), and vagina (Table 1). Strains stored at -80 C were streaked onto tryptic soy agar supplemented with 5% defibrinated sheep blood and incubated anaerobically at 37 °C for 48 h. Subspecies differentiation was performed using a validated qPCR assay targeting subspecies-specific regions of the leukotoxin operon promoter (*lktA*-n for FNN and *lktA*-f for FNF), as previously described (Deters et al., 2024; Kilama et al., 2025). Single colonies were inoculated into pre-reduced, anaerobically sterilized Brain-Heart Infusion broth and overnight cultures were used for DNA extraction.

**Table 1.** Distribution of *Fusobacterium necrophorum* strains utilized in the study by isolation sources, subspecies, clinical vs. non-clinical sources across bovine and human hosts.

| Host | Isolation source | Sample size | *F. necrophorum* subspecies |  |  |  |
| --- | --- | --- | --- | --- | --- | --- |
|  |  |  | *necrophorum* |  | *funduliforme* |  |
|  |  |  | Clinical | Non-clinical | Clinical | Non-clinical |
| Bovine<br>n=84 | Lung | 9 | - | - | - | 9 |
|  | Semen | 25 | - | 1 | - | 24 |
|  | Calf diphtheria | 7 | - | - | 7 | - |
|  | Foot rot | 2 | 1 | - | 1 | - |
|  | Healthy liver | 6 | - | 1 | - | 5 |
|  | Liver abscess | 21 | 9 | - | 12 | - |
|  | Mediastinal Lymph node | 5 | - | - | - | 5 |
|  | Rumen fluid | 8 | - | - | - | 8 |
|  | Vagina | 1 | - | - | - | 1 |
| Human<br>n=53 | Lemierre's syndrome | 11 | - | - | 11 | - |
|  | Tonsillitis | 11 | - | - | 11 | - |
|  | Tonsillar abscess | 5 | - | - | 5 | - |
|  | Other sources* | 26 | - | - | 26 | - |
| **Total** |  | **137** | **10** | **2** | **73** | **52** |

Other sources \* include liver abscess (n=1), liver metastasis (n=1), middle ear granuloma (n=1), peritoneal abscess (n=1), pleural membrane (n=1), septic portal vein thrombosis (n=1), urinary tract proper (n=1), cerebral abscess (n=2), cervical abscess (n=1), colorectal primary tumour (n=1), blood (n=2), cervical lymph node (n=1), liver (n=1), terminal ileum (n=1), abdominal abscess (n=1), blood catheter (n=1), cervical abscess (n=2), urinary tract proper (n=1), and urine (n=1).

### Genomic DNA extraction, library preparation, and whole-genome sequencing

Genomic DNA was extracted from bacterial strains using a modified DNeasy Blood and Tissue Kit protocol (Qiagen, Valencia, CA, USA) optimized for *F. necrophorum*. The bacterial cell pellets were harvested from 800 µL of broth cultures by centrifugation at 20,000 × g for 5 min at 18°C. Cell walls were disrupted using enzymatic lysis buffer containing lysozyme and mutanolysin, followed by incubation at 37°C for 1 h with agitation at 800 rpm. Samples were then treated with 25 µL proteinase K and 200 µL Buffer AL, followed by incubation at 56°C for 30 min. Mechanical cell disruption was achieved by bead beating with 200 mg of sterile 0.1 mm zircon/silica beads for 20 s using a FastPrep-24 bead beater (MP Biomedicals, Irvine, CA, USA). After centrifugation at 13,000 × *g* for 5 min, the supernatant was mixed with 400 µL of 100% ethanol and passed through a DNeasy Mini spin column according to the manufacturer’s instructions. Genomic DNA was eluted in 50 µL of Buffer AE after a 3-5 min incubation at room temperature. DNA purity was assessed using a NanoDrop spectrophotometer (Thermo Fisher Scientific, Waltham, MA, USA), and the DNA was stored at −20°C until library preparation. Whole-genome sequencing was performed by Novogene (Sacramento, CA, USA) using a standardized microbial whole genome sequencing workflow. In brief, genomic DNA libraries were prepared using target organism-specific protocols after sample quality control. Libraries underwent quality control testing before 150 bp paired-end sequencing on the NovaSeq X Plus platform (Illumina Inc., San Diego, CA, USA). Genomic DNA was fragmented, size-selected, end-repaired, A-tailed, and ligated with full-length adapters in accordance with Illumina PE150 protocols.

### Publicly available F. necrophorum strains genome acquisition

Fifty-seven publicly available human *F. necrophorum* genome assemblies were obtained from the National Center for Biotechnology Information (NCBI) Genome database in September 2025. The 57 genomes had high-quality assemblies that met our inclusion criteria. Assemblies were downloaded using the NCBI datasets command-line tool, excluding metagenome-assembled genomes to ensure inclusion of only high-quality isolate genomes. Associated metadata, including accession number, host species, isolation source, geographic origin, and health status, were extracted from NCBI records and manually curated for consistency. The final dataset comprised 137 genomes, representing strains from North America and Europe and encompassing both bovine and human strains across diverse anatomical and clinical contexts. The strains sequenced by our lab (n=80) were classified to subspecies level using a qPCR assay targeting the *lktA*-n (FNN) and *lktA*-f (FNF) genes (Deters et al., 2024), while NCBI genomes were classified based on their subspecies designation in the database. Detailed strain metadata encompassing subspecies identification, host origin, isolation source, and health status for all 137 strains are presented in table 1 and supplementary table S1.

### Genome assembly and annotation

Raw sequencing reads from the 80 strains sequenced by our lab were quality filtered using fastp v0.23.2 (Chen, 2025). Adapter sequences were removed, low-quality bases were trimmed and reads shorter than 100 bp or having a mean Phred quality score below 30 were excluded. Quality-filtered reads were assembled *de novo* using SPAdes v4.2.0 with the “--isolate” flag, which is optimized for bacterial genome assemblies (Prjibelski et al., 2020). Contigs shorter than 500 bp were removed, assembly quality was assessed using QUAST v5.2.0 (Gurevich et al., 2013), and assemblies containing more than 100 contigs were excluded to ensure high-quality genome representation. Genome completeness and contamination were evaluated using CheckM2 v1.1.0 (Chklovski et al., 2024), and assemblies with greater than 5% estimated contamination were removed from further analysis. Taxonomic classification and confirmation were performed using GTDB-Tk v2.5.2 with the Genome Taxonomy Database (GTDB) release 10-RS226 (Parks et al., 2022). To assess the impact of sequencing depth, all samples were subsampled to 100× coverage with Seqtk v1.4 (https://github.com/lh3/seqtk). In each case, the optimal assembly for each sample was selected based on genome completeness, contamination, contig number, and N50. Following assembly, all 137 genomes (80 sequenced + 57 NCBI retrieved) were subjected to comparative genomic analyses. Genome annotation was carried out using Prokka v1.14.6 (Seemann, 2014). Pairwise average nucleotide identity (ANI) was calculated across all 137 genomes using FastANI v1.34 (Jain et al., 2018). An all-versus-all comparison was performed to quantify genome-wide similarity and evaluate clustering patterns within the dataset.

### Pangenome reconstruction and phylogenetic analysis

Pangenome analysis was conducted on all 137 *F*. *necrophorum* assemblies using Panaroo v1.3.4 with strict cleaning parameters to minimize annotation artifacts and enable the automatic removal of invalid gene predictions (Tonkin-Hill et al., 2020). Genes were classified as core (≥ 99%), soft-core (95-99%), shell (15-95%), or cloud (< 15%) based on gene frequency across the dataset. Core gene sequences were aligned using MAFFT v7.526 via Panaroo’s integrated alignment functionality (Katoh and Standley, 2013). Maximum likelihood phylogenetic reconstruction was performed using RAxML-NG v1.2.2 under the GTR+G substitution model with 1,000 bootstrap replicates (Kozlov et al., 2019). The tree was visualized using iTOL v6.8.1 (Letunic and Bork, 2024), incorporating metadata layers for subspecies, ecological lifestyle, host origin, sample type, disease status and geographic location.

### Functional profiling

Predicted proteins from each genome (Prokka-generated faa files) were aligned to the KEGG prokaryotic protein database (release 2025-09-21) using DIAMOND BLASTP v2.2.15 with an e-value cutoff of 1 × 10□¹□. KEGG orthology (KO) assignments were derived from significant alignments and filtered using minimum amino acid identity (≥40%) and query coverage (≥80%) thresholds. Unique KOs were retained for each genome for downstream analyses. To reconstruct metabolic pathway potential, genome-specific KO sets were analyzed using MinPath v1.6 (Ye and Doak, 2009) to infer parsimonious KEGG pathway complements. MinPath outputs were compiled into pathway presence/absence matrices across all genomes.

### Virulence factor and CRISPR-Cas system detection

Virulence factors were identified by aligning predicted protein sequences against a curated *F. necrophorum*-specific virulence database (Carrara et al., 2024), using DIAMOND BLASTP v2.2.15 (Buchfink et al., 2021). CRISPR-Cas systems were identified by searching predicted protein sequences against curated Cas protein sequences from UniProt (Consortium, 2025) using the same approach. Hits were retained using minimum thresholds of ≥80% amino acid identity and ≥80% query coverage for virulence factors, and ≥70% amino acid identity and ≥60% query coverage for Cas proteins.

### Antimicrobial resistance genes and plasmid identification

The Resistance Gene Identifier (RGI) v6.0.3 (Alcock et al., 2020) was used to screen genomes against the Comprehensive Antibiotic Resistance Database (CARD) (McArthur et al., 2013) to identify ARGs. Only “strict” and “perfect” hits were retained. Proksee (build/version 2026-02-25) (Grant et al., 2023) was used to visualize the genomic arrangement of ARGs when multiple ARGs were co-located on a single contig. MOB-suite v3.1.9 (Robertson et al., 2020) was used to identify and characterize plasmid sequences from the genome assemblies. PlasmidFinder v2.1.6 (Carattoli et al., 2014) was also used with default parameters and the integrated plasmid database to confirm plasmid sequence detection.

### Statistical analysis

All statistical analyses were performed in R v4.5.2 (R Core Team, 2025) using the packages *dplyr, ggplot2*, and *ComplexHeatmap*. Statistical significance was defined as α = 0.05 for all tests. Comparative genomics analyses employed non-parametric Mann-Whitney U tests for pairwise comparisons of genomic features (genome size, G+C content, CDS count, CRISPR-Cas-associated genes) between groups, as normality assumptions were violated based on Shapiro-Wilk tests. Fisher’s exact test was used to compare the prevalence of virulence factor genes and ARGs between groups.

For virulence factor categories, samples were classified as positive using an “any-variant” approach (positive if any variant within the category was detected). Odds ratios with 95% confidence intervals were calculated for significant associations. Multiple testing correction was applied using the Benjamini-Hochberg false discovery rate (FDR) method. Effect sizes were assessed using Cliff’s delta for continuous variables and Cramér’s V for categorical variables. Data visualization included hierarchical clustering using Euclidean distance with complete linkage for heatmap construction (Gu et al., 2016). Additional visualizations were generated using GraphPad Prism v10 (GraphPad Software, San Diego, CA, USA).

## RESULTS

### Summary of strains origin and subspecies distribution

A total of 137 genomes were included in this study, comprising strains sequenced in this study (n = 80) and publicly available genomes retrieved from NCBI (n = 57) (Supplementary Table S1). The FNN strains (n = 12) were recovered exclusively from beef cattle, predominantly from liver abscesses (n = 9), with additional strains from semen (n = 1), healthy liver tissue (n = 1), and foot rot (n = 1). However, 125 FNF genomes were derived from both cattle (n = 72) and humans (n = 53), spanning a wide range of non-clinical and disease-associated niches. Bovine FNF strains were largely isolated from bull semen (n = 24), liver abscesses (n = 12), lungs (n = 9), ruminal fluids (n = 8), and diphtheretic lesions (n = 7) (Supplementary Table S1). Human FNF genomes originated from patients with tonsillitis (n = 15) and Lemierre’s syndrome (n = 11), as well as other clinical samples such as tonsillar samples (n = 5), cervical abscesses (n = 3), and cerebral abscesses (n = 2). Although all human strains were mainly associated with clinical conditions, bovine strains were sourced from both healthy or non-clinical (n = 54) and clinical conditions (n = 30), providing a broad representation of both commensal and clinical lifestyles (Table 1). Geographically, the genomes were mainly originated from North America and Europe, with the majority recovered from the U.S. (n = 85). Additional genomes were originated from Switzerland (n = 27), Denmark (n = 22), Spain (n = 2), and Canada (n = 1). Genomes of FNF were distributed across all represented countries, whereas FNN genomes were predominantly of U.S. origin (n = 11), with a single isolate originated from Switzerland (n = 1).

### Genome assembly, annotation, pangenome and comparative genome analysis

Of the 80 *F. necrophorum* genomes sequenced in the present study, mean sequencing coverage was 174 ± 4.5× (standard error of the mean [SEM]), with a genome size of 2,216,055 ± 22,049 bp and G+C content of 34.7 ± 0.1%. The number of coding sequences per genome was 2,060 ± 22.6 (Table S2, Fig. S2). These genomes contained 45 ± 0.2 tRNA genes, 2-9 rRNA genes, and 877 ± 18.5 hypothetical protein-coding regions on average. CRISPR-Cas proteins were identified in all 80 genomes, with an average of 3.3 ± 0.1 per genome (Table S2, Fig. S2).

The pangenome comprised 5,820 predicted genes (Fig. 1A), including 1,495 core genes present in ≥ 99% of genomes and 33 soft core genes. The accessory genome (cloud + shell) comprised 4,292 genes (Fig. 1A). The size of the accessory genome and the gene accumulation curve indicated that *F. necrophorum* possesses an open pangenome (Fig. 1B, C).

**Figure 1.**
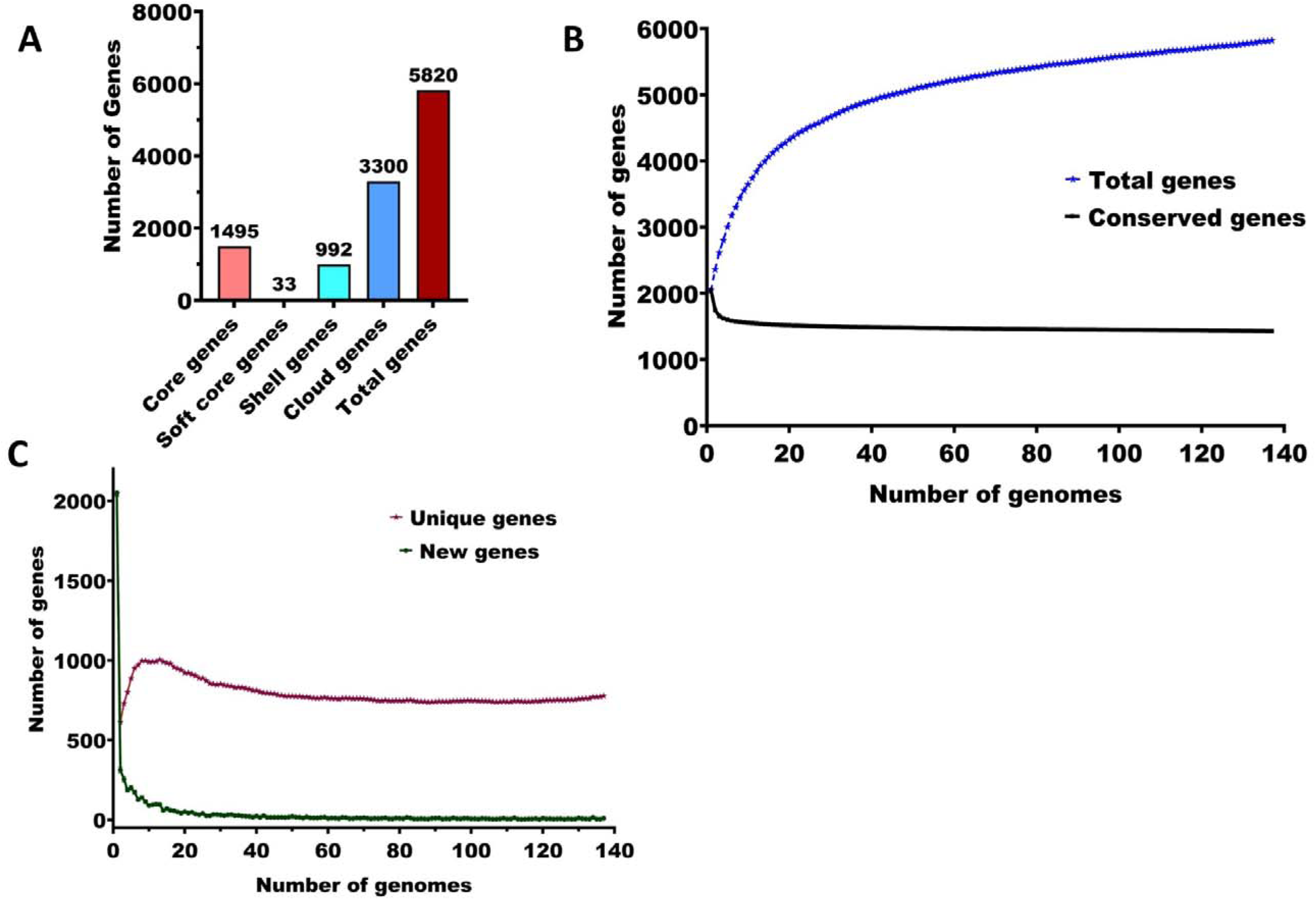
Pangenome analysis of 137 *Fusobacterium necrophorum* genomes. **(A)** Gene category distribution showing absolute counts of core genes (present in 99-100% of genomes, n=1,495), soft core genes (95-99%, n=33), shell genes (15-95%, n = 992), and cloud genes (0-15%, n=3,300). **(B)** Pangenome accumulation curve showing the increase in total gene count with sequential genome addition, while the number of conserved genes stabilizes at approximately 1,400 genes. **(C)** Gene discovery dynamics illustrating the decline in newly identified genes with increasing number of genome and the cumulative number of unique genes approaching stabilization at approximately ∼700 genes.

### Phylogenetic analysis and average nucleotide identity

Phylogenetic reconstruction based on core genome alignment revealed clear subspecies-level structure among the 137 genomes (Fig. 2). The phylogeny resolved two major clades corresponding to the recognized subspecies FNF and FNN, with strong bootstrap support. Average nucleotide identity across all genomes ranged from 95.6% to 100%, confirming all strains as *F. necrophorum* (≥ 95% species threshold). Intra-subspecies comparisons showed high similarity (FNF: mean 98.6%, range 95.9–100%; FNN: mean 99.4%, range 98.5–100%), whereas inter-subspecies comparisons exhibited lower mean ANI (97.1%, range 95.62–99.9%). Bovine and human strains were largely clustered separately within subspecies clades.

**Figure 2:**
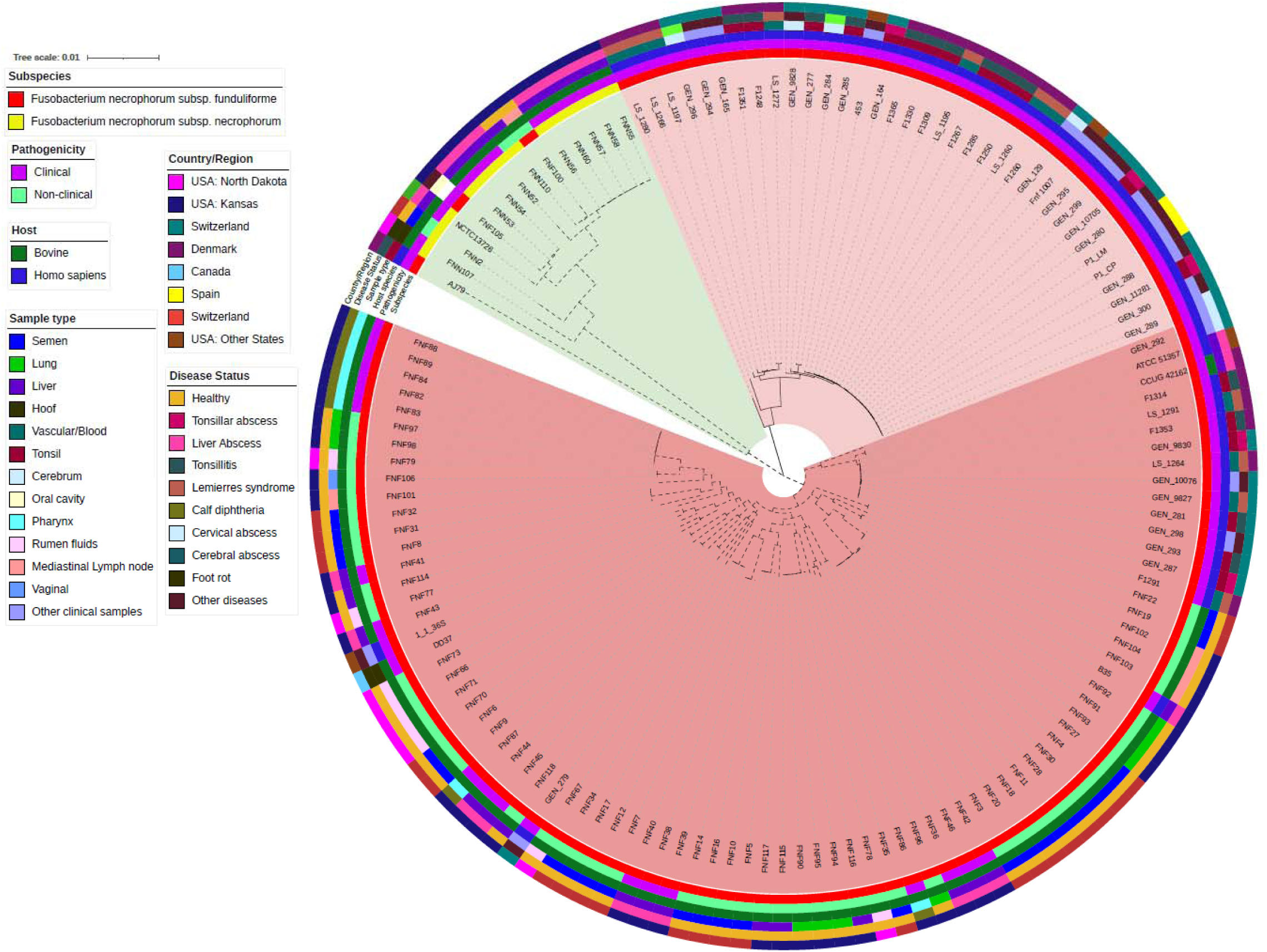
Maximum likelihood phylogeny of *Fusobacterium necrophorum* strains. The tree was inferred from an alignment of 1,495 core genes identified through pangenome analysis, including strains from this study (n = 80) and and publicly available genomes (n = 57). The horizontal scale bar represents substitutions per nucleotide. Rings from outside to inside represent country of strains origin, disease status, sample type, host species, ecological lifestyle, and subspecies classification. Inner shading denotes subspecies: light green for FNN and two shades of red representing the two FNF clades.

There was slightly higher genomic similarity within the host groups based on ANI (bovine-bovine: 98.5%; human–human: 99.0%) than between hosts (bovine–human: 98.2%) (Table S3). Certain *F. necrophorum* strains recovered from different host species appeared to be nearly identical and may represent the same strain (ANI >99.9%). Notable cross-host examples included ATCC 51357 (Human; oral sample; USA) and CCUG 42162 (bovine; liver abscess; Denmark), LS_1291 (human; Lemierre’s syndrome; Denmark) and CCUG 42162 (bovine; liver abscess; Denmark), and F1314 (human; tonsillitis; Denmark) and CCUG 42162 (bovine; liver abscess) (Supplementary Table S3). Pathogenic strains (clinical; n = 83) and non-clinical strains (healthy; n = 54) were distributed in both subspecies’ clades with no distinct clustering by health status.

Bull seminal strains, which represented majority of non-clinical *F. necrophorum* strains, demonstrated high genetic conservation, characterized by tight phylogenetic clusters with high intra-group ANI values. Representative pairs included FNF34 and FNF17 (99.9% ANI), FNF3 and FNF20 (99.9% ANI), and 45 additional intra-group comparisons among 23 bull seminal strains showing consistently elevated ANI (Supplementary Table S3). Human clinical strains showed limited disease-specific phylogenetic clustering. Strains from patients with tonsillitis, Lemierre’s syndrome, and various abscess-associated conditions were distributed across FNF clades, primarily according to subspecies classification rather than clinical presentation. Geographic origin did not appear to drive phylogenetic clustering, as strains from the USA, Denmark, Switzerland, and other countries were distributed throughout subspecies clades (Fig. 2).

### Functional genomic profiling

#### Functional differences between FNN and FNF strains

Functional profiling was performed to characterize differences in gene content between the two FN subspecies. We observed significant differences in KO (KEGG Orthology) prevalence between *F. necrophorum* subspecies, with 80 KOs differing significantly between FNF (n = 125) and FNN (n = 12) strains (FDR < 0.05; Fisher’s exact test) (Fig. 3A). The FNN strains showed significantly higher prevalence of KOs associated with carbohydrate transport and metabolism, including raffinose/stachyose transport systems (e.g., *msmE*, K10117; *msmF*, K10118; *msmG*, K10119), which were present in 100% of FNN strains but only 0.8% of FNF strains (p < 0.001). The KOs involved in advanced glycation end-product (AGE) degradation pathways (e.g., *frlB,* K10708*; frlD,* K10710, fructoselysine metabolism) were also specific to FNN strains (100% vs 0% in FNF).

**Figure 3.**
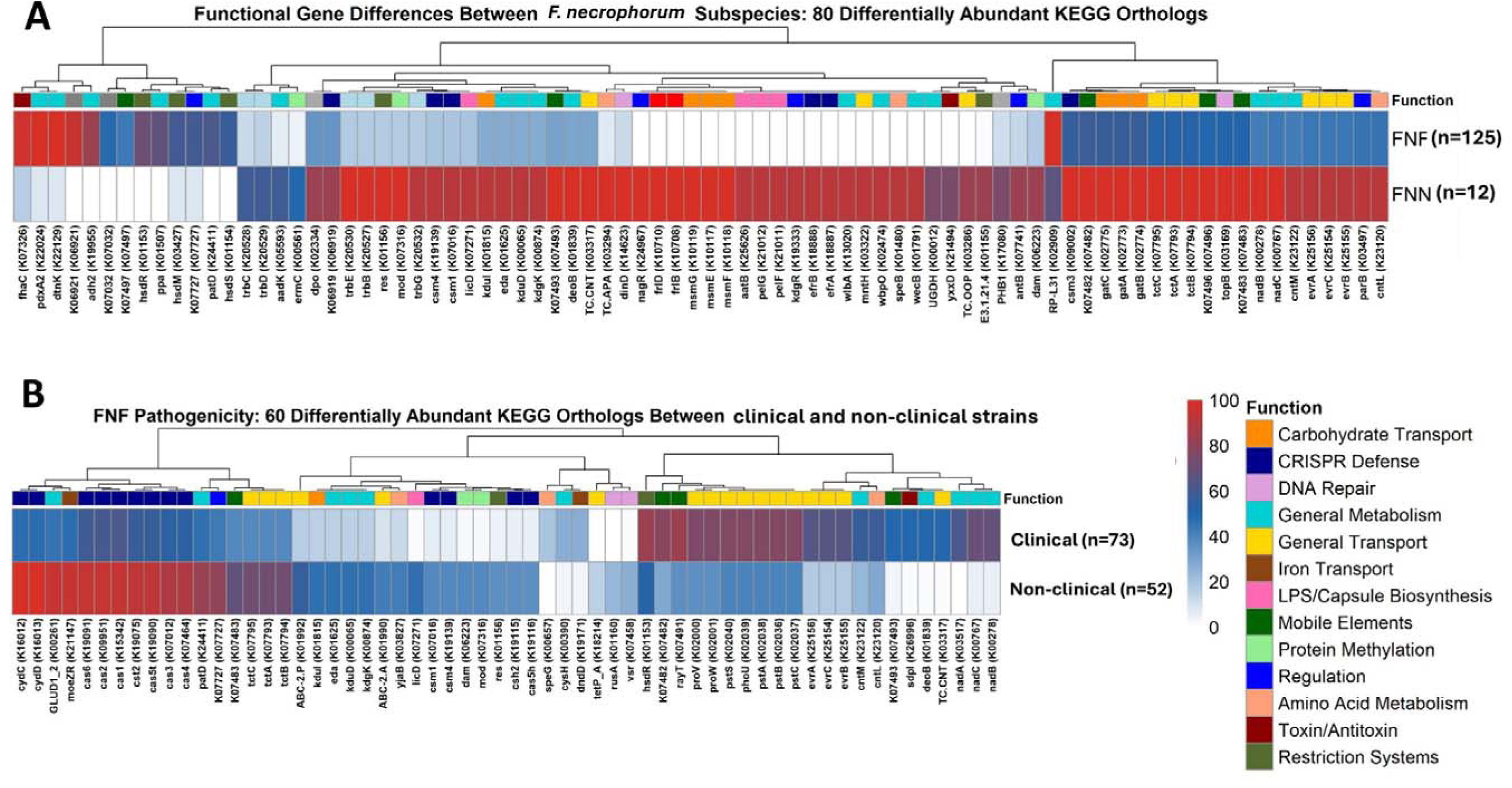
Functional differences in *Fusobacterium necrophorum* reveal subspecies specialization and pathogenic adaptations. Heatmaps showing the prevalence (%) of KEGG orthologs (KOs) that differ significantly by subspecies and by pathogenicity. (A) Subspecies comparison showing 80 differentially abundant functions between FNF (n=125) and FNN (n=12) strains, highlighting FNF specialization in hemolysin systems and threonate metabolism versus FNN carbohydrate transport capabilities. (B) Pathogenicity analysis revealing 60 differentially abundant functions between pathogenic and non-clinical FNF strains across both human and bovine hosts. Functions are clustered by similarity and color-coded by predicted biological roles, highlighting subspecies-specific metabolic specializations. Statistical significance: FDR < 0.05, Fisher’s exact test. Color scale represents prevalence (%): white denotes 0%, blue denotes intermediate (∼50%), and red denotes 100%, with color intensity proportional to prevalence values.

In contrast, FNF strains showed higher prevalence of KOs related to threonate metabolism and hemolysin-associated systems. Nearly all FNF strains (99.2%) contained genes assigned to KOs corresponding to D-threonate/D-erythronate metabolism (e.g *pdxA2*, K22024; *dtnK*, K22129), which were rare in FNN strains (8.3%, *P* < 0.001). The FNF strains also had a higher prevalence of KOs associated with hemolysin activation (e.g., *fhaC,* K07326) (97.6% vs 16.7%, *P* < 0.001), type I restriction-modification (e.g., *hsdR,* K01153*; hsdS,* K01154) (56.0-70.4% vs 0%, *P* < 0.001), and alcohol dehydrogenase (e.g., *adh2,* K19955) (84.0% vs 0%, *P* < 0.001). Overall, these findings indicate distinct metabolic and functional specialization between FNN and FNF, reflecting their adaptation to different ecological niches.

#### Functional differences between non-clinical and clinical FNF strains independent of host origin

Comparison of clinical (n = 73) and non-clinical FNF (n = 52) strains identified 60 KOs that differed significantly in prevalence (FDR < 0.05, Fisher’s exact test) (Fig. 3B). Non-clinical FNF strains showed near-universal presence of KOs associated with cytochrome oxidase components (e.g., *cydC,* K16012*; cydD,* K16013) (100% vs 47.9%, *P* < 0.001), which are essential for aerobic respiration under microaerobic conditions. Non-clinical FNF strains were also enriched for KOs associated with CRISPR-Cas (e.g., *cas6,* K19091) (94.2% vs 58.9%, *P* < 0.001), DNA methylation systems (e.g., *dam,* K06223*; mod,* K07316) (38.5% vs 2.7%, *P* < 0.001), and type III restriction enzymes (36.5% vs 4.1%, *P* < 0.001). On the other hand, clinical FNF strains showed higher prevalence of KOs involved in glutamate metabolism (e.g., *gdhA,* K00261) (94.2% vs 45.2%, *P* < 0.001) and specialized transport systems compared to non-clinical strains.

#### Functional differences between bovine and human clinical FNF strains

A total of 52 KOs were different in prevalence between clinical FNF strains from bovine (n = 20) and human (n = 53) hosts (FDR < 0.05, Fisher’s exact test) (Supplementary Table S4). Clinical bovine FNF strains had a higher prevalence of KOs associated with respiratory and metabolic processes compared to human FNF strains. All bovine FNF strains contained KOs associated with glutamate dehydrogenase (e.g., *gdhA,* K00261) (100% vs 24.5% in human strains, *P* < 0.001) and cytochrome bd oxidase components (e.g., *cydC,* K16012; *cydD,* K16013) (100% vs 28.3%, *P* < 0.001). Bovine strains were also enriched for KOs related to carbohydrate catabolism, including 2-dehydro-3-deoxy-D-gluconate metabolism (e.g., *kduD,* K00065; *kdgK,* K00874; *eda,* K01625; *kduI*, K01815) (50% vs 1.9%, *P* < 0.001) and ABC-2 type transporters (30-60% vs 0-1.9%, *P* < 0.001) as compared to strains from humans. The KOs associated with CRISPR-Cas (e.g., *cas6,* K19091; *cas2,* K09951; *cmr4,* K09000; *cmr6,* K19142) were likewise more prevalent in bovine clinical FNF strains (25-90% vs 0-49.1%, *P* < 0.05) than those from humans.

However, clinical human FNF strains exhibited preferential enrichment of KOs related to nicotinamide adenine dinucleotide (NAD) biosynthesis and nucleoside metabolism. The KOs corresponding to NAD biosynthesis (e.g., *nadA,* K03517; *nadB,* K00278; *nadC,* K00767) were detected in 83.0-92.5% of human strains compared to 5-10% of bovine strains (*P* < 0.001). Human FNF strains also showed higher prevalence of KOs associated with DNA replication initiation (*dnaC*, K02315) (92.5% vs 20%, p < 0.001), nucleoside transport (*deoB,* K01839) (67.9% vs 0%, *P* < 0.001), and glycine betaine/proline transport (*proV*, K02000) (92.5% vs 30%, *P* < 0.001).

#### Functional differences between bovine clinical and non-clinical bovine FNF strains

Analysis of bovine FNF strains revealed minimal differences in KO prevalence between clinical (n = 52) and non-clinical (n = 20) strains. Among the 1,153 KOs identified only one KO that differed significantly in prevalence (FDR < 0.05, Fisher’s exact test). The KO corresponding to inorganic pyrophosphatase (*ppa*, K01507) was more prevalent in non-clinical strains than in clinical strains (80.8% vs 25.0%, *P* = 0.020) (Supplementary Table S5).

#### Functional differences in genomes of bovine anatomical site specific FNF strains

Pairwise comparison of FNF strains from seven bovine anatomical sites or conditions identified site-specific differences in KO prevalence. Of 21 possible comparisons, four showed significant differences (FDR < 0.05, Fisher’s exact test) between different anatomic sites. The seminal strains differed from other anatomical sites, with differential prevalence of 9 KOs compared to calf diphtheria-associated strains, 8 compared to lung strains, and 5 compared to mediastinal lymph node-associated FNF strains. Liver abscess-associated strains also differed from calf diphtheria-associated strains in 4 KOs (Supplementary Table S6). Calf diphtheria-associated strains exhibited higher prevalence of KOs corresponding to tetracycline resistance systems [e.g., *tet*(M), K18829; *tet*(O), K18830], transcriptional regulators (e.g., *rutR,* K09017), and beta-exotoxin transport systems (K25147, K25148) that were absent (0% prevalence) in semen and liver abscess strains. Similar to calf diphtheria-associated strains, lung and mediastinal lymph node strains demonstrated high prevalence of tetracycline resistance and beta-exotoxin transport KOs when compared to seminal strains. The lung-associated strains also showed enrichment of DNA recombination genes (e.g., *rusA*, K01160; *recT*, K07455) and type I restriction enzymes (e.g., *hsdR,* K01153) (Supplementary Table S6). The seminal FNF strains displayed distinct KO prevalence patterns, including KOs corresponding to chromosome partitioning proteins (e.g., *parB,* K03497) (83.3% vs 0% in calf diphtheria), mobile genetic elements (K07483 transposase) (83.3% vs 11.1% in lung strains), and inorganic pyrophosphatase (*ppa,* K01507) (95.8% vs 0% in calf diphtheria). Calf diphtheria-derived FNF strains were additionally enriched for KOs associated with DNA modification systems, including 23S rRNA methyltransferase (*rlmA1*, K00563) and restriction enzymes (*mcrA,* K07451) (71.4% vs 0% in semen strains).

### Virulence factor repertoire

#### Distribution of virulence factors in F. necrophorum genomes

Using virulence factor database (VFDB) and a curated *F. necrophorum*-specific database, 84 unique virulence factor protein variants were identified (Supplementary Table S7). These variants were grouped into 13 functional categories and showed heterogeneous distribution across anatomical sites in bovine strains (Fig. 4A) and disease conditions in both bovine and human hosts (Fig. 4B). Identified categories included autotransporter/outer membrane proteins, hemagglutinin adhesins, leukotoxins, LPS/LOS biosynthesis enzymes, CRISPR-associated proteins, virulence-associated proteins, and toxin-related factors (Supplementary Fig. S3). Core virulence factors (≥95% of genomes; 27 variants, 32.1%) were primarily composed of LPS/LOS biosynthesis enzymes (7/8 variants), leukotoxin components (6/13 variants), and hemagglutinin proteins (5/14 variants). Accessory virulence factors (15-95% prevalence; 30 variants, 35.7%) included autotransporter proteins (7/14 variants), leukotoxin variants (6/13 variants), and CRISPR-associated proteins (5/8 variants). Rare virulence factors (<15% prevalence; 27 variants, 32.1%) were dominated by autotransporter proteins (7/14 variants), hemagglutinin variants (6/14 variants), and virulence-associated proteins (5/8 variants) (Supplementary Table S7).

**Figure 4.**
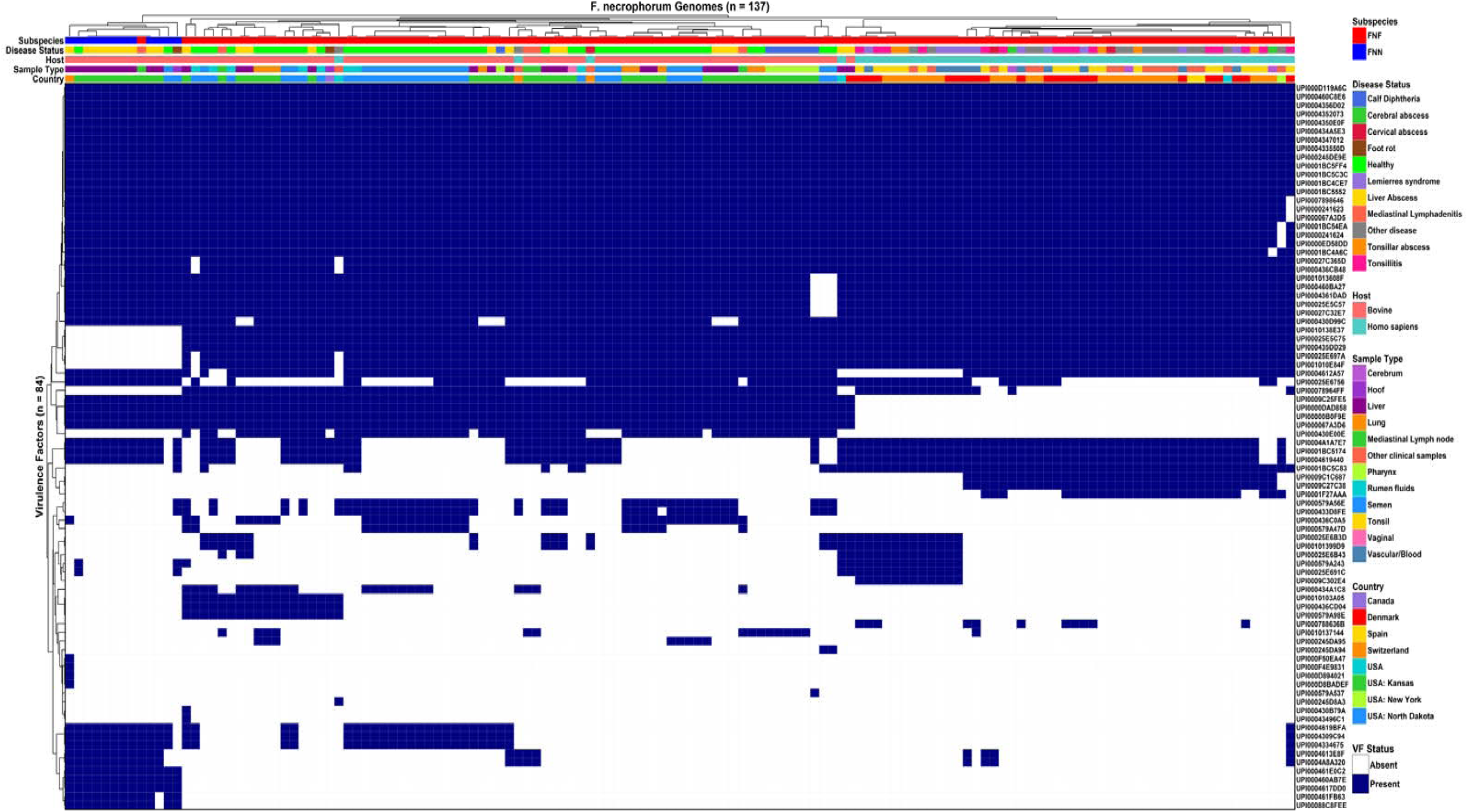
Heatmap showing presence/absence of 84 virulence factors detected across 137 *Fusobacterium necrophorum* genomes at ≥80% identity and ≥80% coverage using a custom *Fusobacterium* database plus VFDB. Top annotation bars indicate subspecies (*Fusobacterium necrophorum* subsp. *funduliforme* [FNF]*; Fusobacterium necrophorum* subsp. *necrophorum* [FNN]), disease status, host species, sample type, and geographic origin. Blue indicates presence and white indicates absence. Hierarchical clustering demonstrates subspecies-specific and disease-associated patterns of virulence factor distribution.

#### Comparison of virulence factors between bovine FNF and FNN strains

Comparative analysis of bovine FN strains revealed subspecies-specific differences in invasion-related virulence factors alongside conservation of core pathogenicity determinants (Table 2). Five virulence factors differed significantly between FNF (n = 72) and FNN (n = 12) strains (FDR < 0.001). The FNN strains exhibited exclusive presence of invasion-associated factors, including a *FadA* adhesion protein (UPI000460AB7E), a GDSL lipase/esterase (UPI0004617DD0), and a serine protease (UPI000461E0C2), each detected in 100% of FNN strains but only 1.4% of FNF strains (FDR = 1.61×10□¹²). Autotransporter outer membrane proteins (UPI00025E5C75, UPI000435DD29) were present in 98.6% of FNF strains but absent in all FNN strains. However, 24 virulence factors were universally conserved across both subspecies, including multiple leukotoxin-associated proteins (including *lktA, lktB* and *lktC),* LPS/LOS biosynthesis enzymes, and immune evasion-related proteins.

**Table 2.**
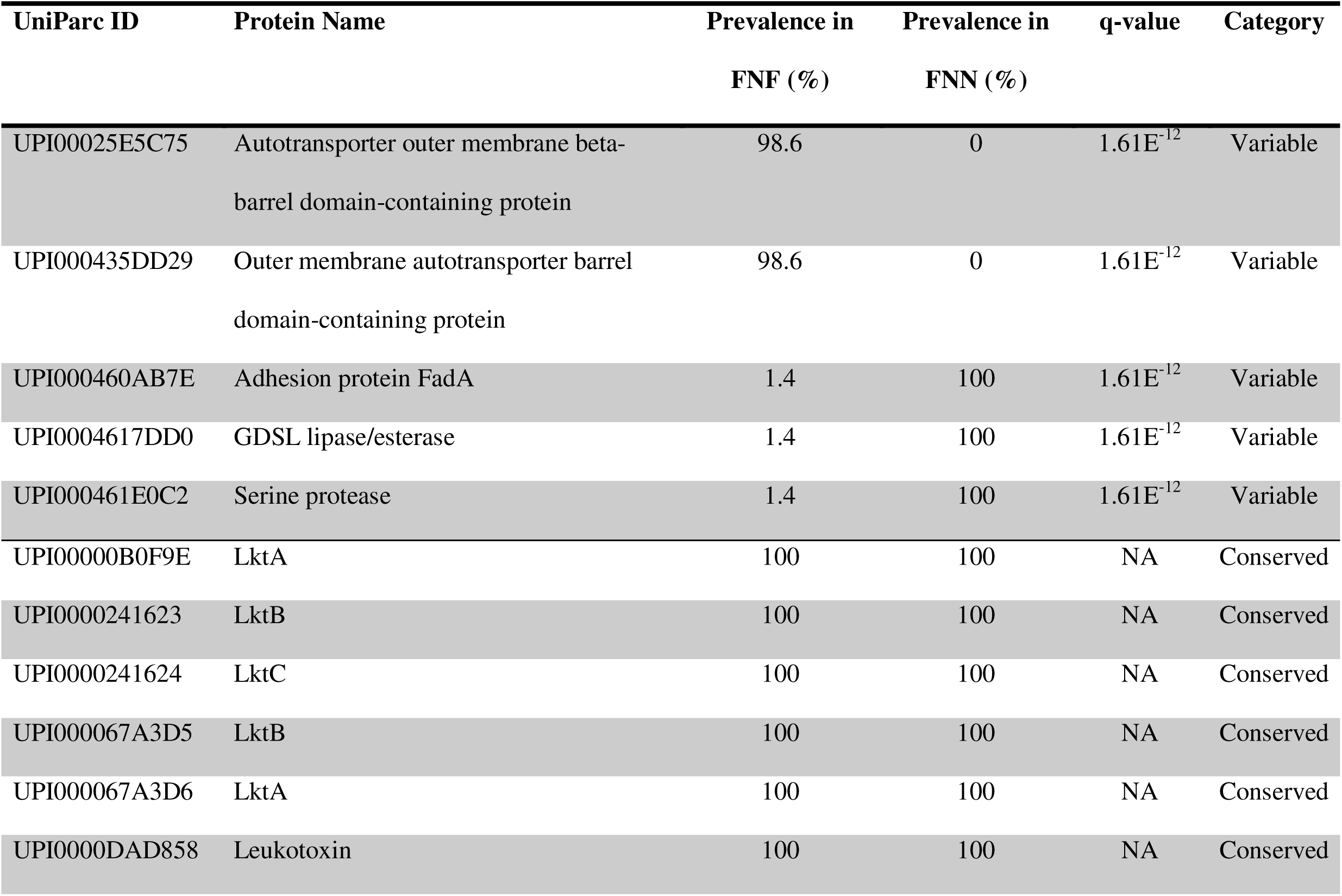
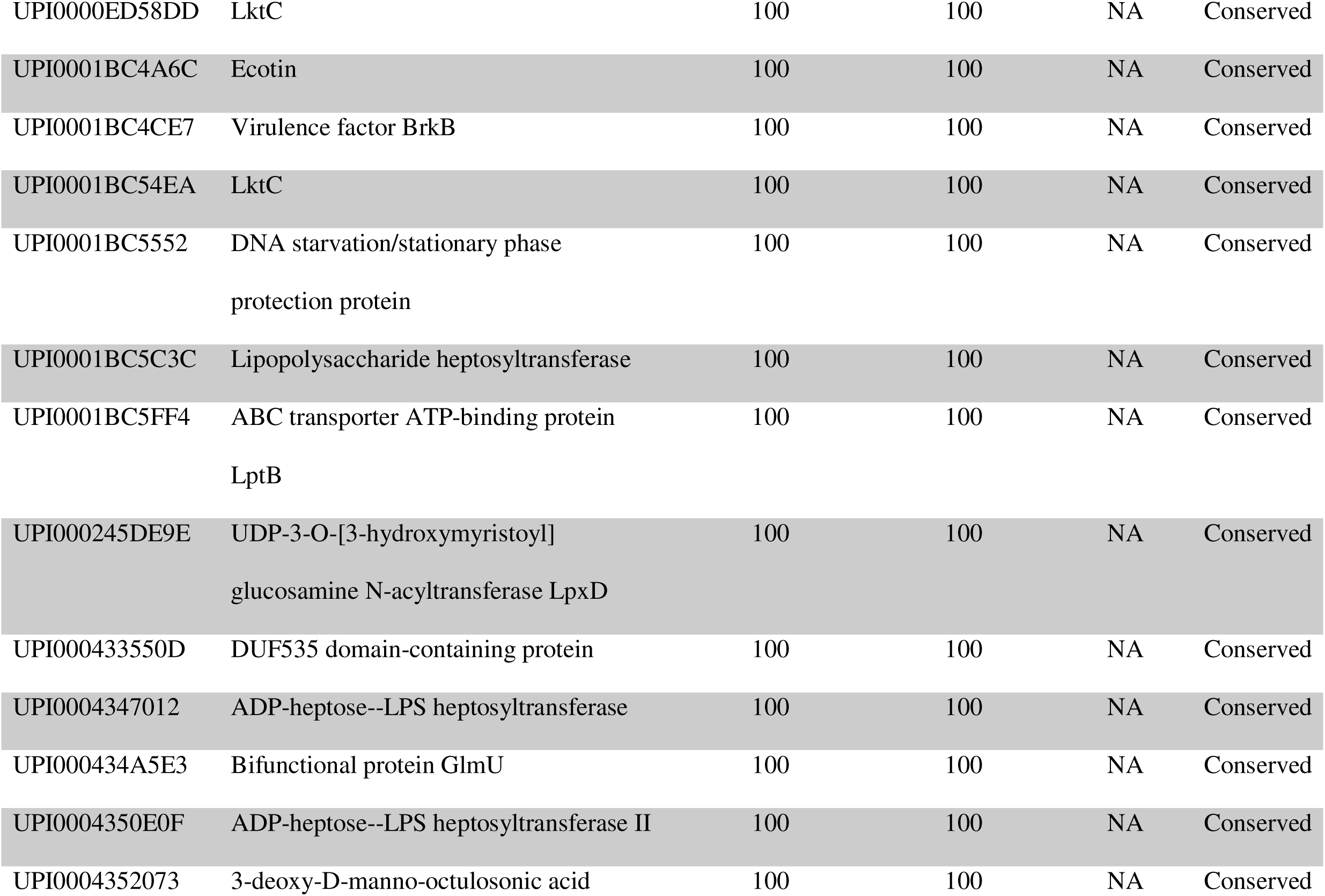
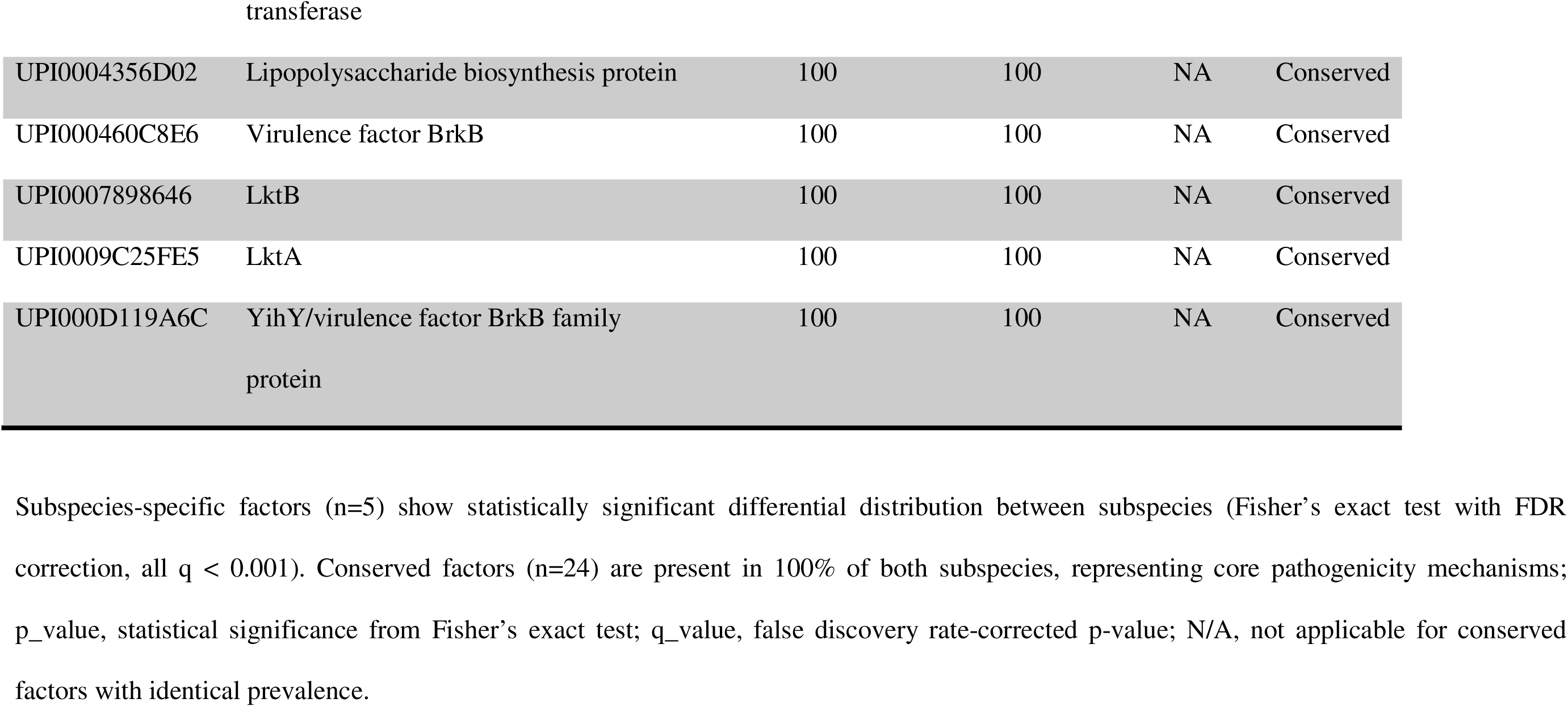
Subspecies-specific and conserved virulence factor distribution in bovine *Fusobacterium necrophorum* strains.

#### Differences in virulence factors between bovine and human clinical and non-clinical FNF strains

Comparison of clinical (n=73) and non-clinical (n=52) FNF strains revealed distinct lifestyle-associated virulence patterns, with 27 virulence factors differing significantly in prevalence (FDR < 0.05) (Fig. 5). The virulence factors (n=15) in non-clinical strains were predominantly associated with adhesion and host interaction functions. Multiple leukotoxin variants (including *lktA* variants UPI00000B0F9E, UPI000067A3D6, UPI0009C25FE5 and leukotoxin UPI0000DAD858) were present in 100% of non-clinical FNF strains compared to 32.9% of clinical FNF strains (FDR = 1.60×10□¹□). Filamentous hemagglutinin proteins and CRISPR-associated endoribonucleases were also more prevalent in non-clinical FNF strains (36.5-82.7% vs 1.4-27.4%). In contrast, pathogenicity-associated virulence factors (n=12) predominantly encoded for toxin-mediated and membrane-associated functions. Fic family toxin-antitoxin systems (UPI0001BC5174, UPI0004619440, UPI0004A1A7E7) were present in 78.1% of clinical strains compared to 36.5% of non-clinical FNF strains (FDR = 2.75×10□□). Alternative *lktA* variants showed exclusive association with pathogenic strains (47.9% vs 0%; FDR = 1.19×10□□), along with enrichment of autotransporter proteins and metabolic enzymes.

**Figure 5.**
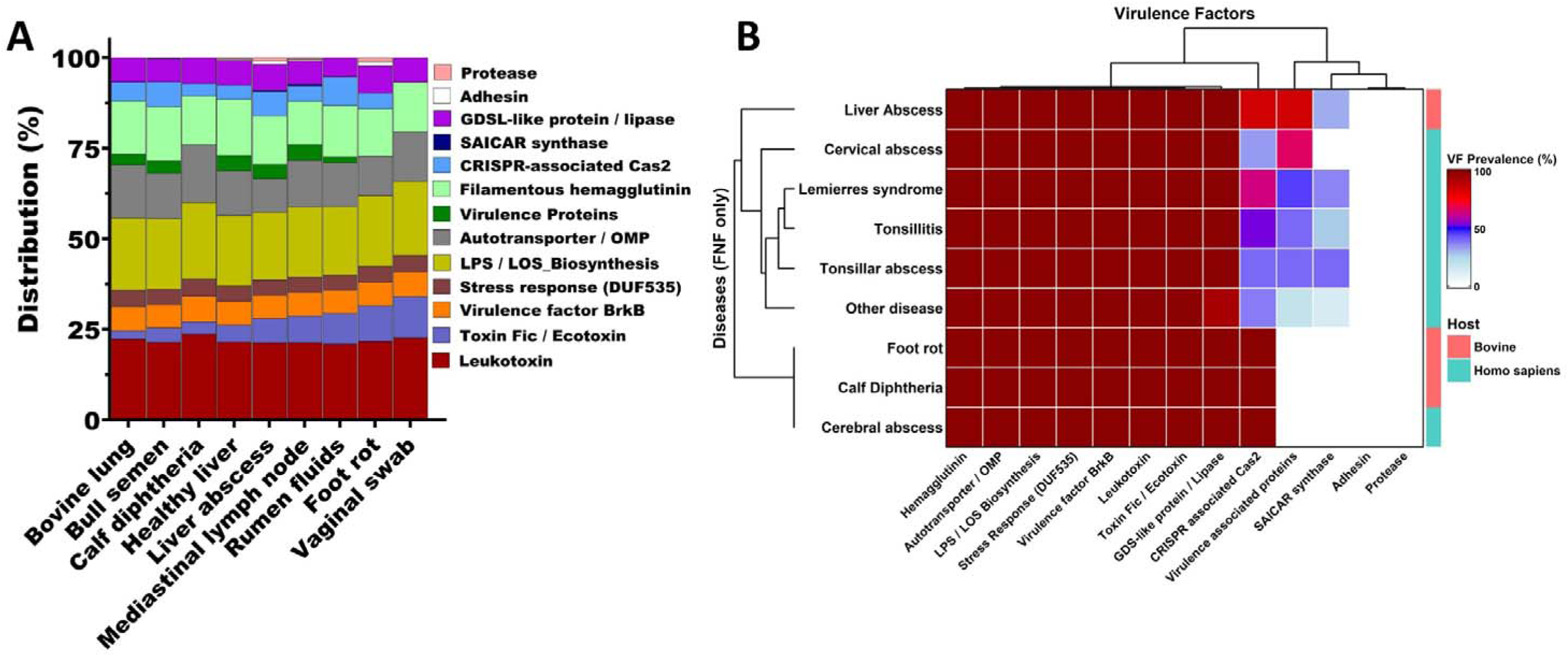
Differential distribution of virulence factors (VF) between pathogenic and non-clinical *Fusobacterium necrophorum* subsp. *funduliforme* strains. Heatmap showing the prevalence of 27 VFs significantly associated with either clinical (diseased, n=73) or with non-clinical (healthy, n = 52) strains. Columns represent individual VFs (UniParc IDs), and rows represent isolate groups. The top annotation bar indicates functional categories. Color intensity (blue-red-white) reflects prevalence (0–100%). A total of 12 virulence factors were enriched in pathogenic strains, while 15 were enriched in non-clinical strains. All differences were statistically significant (Fisher’s exact test with FDR correction, q < 0.05).

#### Differences in virulence factors between bovine and human FNF strains

Host-specific analysis revealed significant differences in virulence factor distribution between bovine (n=72) and human (n=53) FNF strains (Table 2). A total of 31 virulence factors differed significantly between human and bovine FNF strains (FDR < 0.05). Bovine FNF strains were characterized by near-universal presence of multiple leukotoxin variants, with four *lktA* variants and one leukotoxin variant detected in 100% of bovine strains but only 7.5% of human strains (FDR = 3.75×10□²□). Filamentous hemagglutinin proteins (84.7% vs 3.8%; FDR = 2.02×10^□²□^), along with autotransporter proteins (97.2% vs 35.8%; FDR = 9.40×10□¹□) were also strongly enriched in bovine strains. The human-derived strains were characterized by higher prevalence of toxin-associated systems, particularly Fic family toxin-antitoxin systems (94.3% vs 12.5%; FDR = 1.40×10□²□), as well as alternative leukotoxin variants (66% vs 0%; FDR = 5.53×10□¹□).

#### Differences in virulence factors between bovine clinical and non-clinical FNF strains

Within bovine FNF strains, differences in virulence factor prevalence between clinical (n=20) and non-clinical (n = 52) strains were limited. Of 84 identified virulence factors, 24 were present in both and 50 were variable among clinical and non-clinical FNF strains. Only one virulence factor, LOS biosynthesis enzyme (UPI000430D99C), showed a significant difference (FDR = 0.017), with higher prevalence in non-clinical than clinical strains (98.1% vs 65.0%). This association was not observed in the multi-host analysis (Supplementary Table S8), indicating a cattle-specific effect.

#### Comparison of virulence factors in FNF strains across different bovine anatomical sites

There were 10 virulence factors that differed significantly in prevalence across strains from different anatomical sites (FDR < 0.05) (Supplementary Table S9). An autotransporter outer membrane protein (UPI0010137144) was enriched in mediastinal lymph node (4/5, 80%), laryngeal (calf diphtheria;5/7, 71.4%), and lung strains (5/9, 55.6%), but absent in strains from semen, liver abscesses, rumen fluid, and healthy liver tissue (FDR = 8.5×10□□). Virulence-associated protein E variants (UPI0004613E8F, UPI0004A8A320) were prevalent 3 of 5 mediastinal lymph node strains and rare or absent in other anatomical sites (FDR = 4.8×10□□). Fic toxin systems (UPI0001BC5174, UPI0004619440, UPI0004A1A7E7) showed high site-specific enrichment in ruminal fluid (7/8, 87.5%) and mediastinal lymph node (4/5, 80%) but were less frequent in semen (6/24, 25%) and absent in lung (0/9) strains. Filamentous hemagglutinin proteins (UPI000433D8FE, UPI000579A56E) were most prevalent in the genomes of healthy liver (5/5, 100%) and seminal strains (19/24, 79.2%) and were reduced or absent in laryngeal- (calf diphtheria;1/7, 14.3%) and mediastinal lymph node-origin strains (0/5). The lipooligosaccharide (LOS) biosynthesis enzyme (UPI000430D99C) was present in all strains originated from semen, rumen fluid, healthy liver, and mediastinal lymph node, but had low prevalence in liver abscess-associated (6/12, 50%) strains.

### Antimicrobial resistance gene profiles

Six ARGs were identified across the 137 *F. necrophorum* genomes (Table 3). Most genomes (n = 106) did not carry any ARGs. The ribosomal protection protein gene *tet*(O), conferring resistance to tetracycline, was the most prevalent ARG (21.2% of genomes), followed by *erm*(B) (8.8%), *tet*(40) (7.3%), *tet*(M) (1.5%), *tet*(36) (0.7%), and *estT* (0.7%) (Table 3). Collectively, the ARGs identified conferred resistance to two antimicrobial classes: tetracyclines (22.6%) and macrolides (8.8%). Among bovine *F. necrophorum* strains, ARG carriage was higher in FNN than in FNF (58.3% vs 29.2%), although this difference was not statistically significant (*P* = 0.094) (Fig. 6A). However, *erm*(B) (50.0% vs 8.3%, *P* = 0.001) and *tet*(O) (58.3% vs 27.8%, *P* = 0.048) were significantly more prevalent in FNN than FNF strains (Fig. 6B).

**Figure 6.**
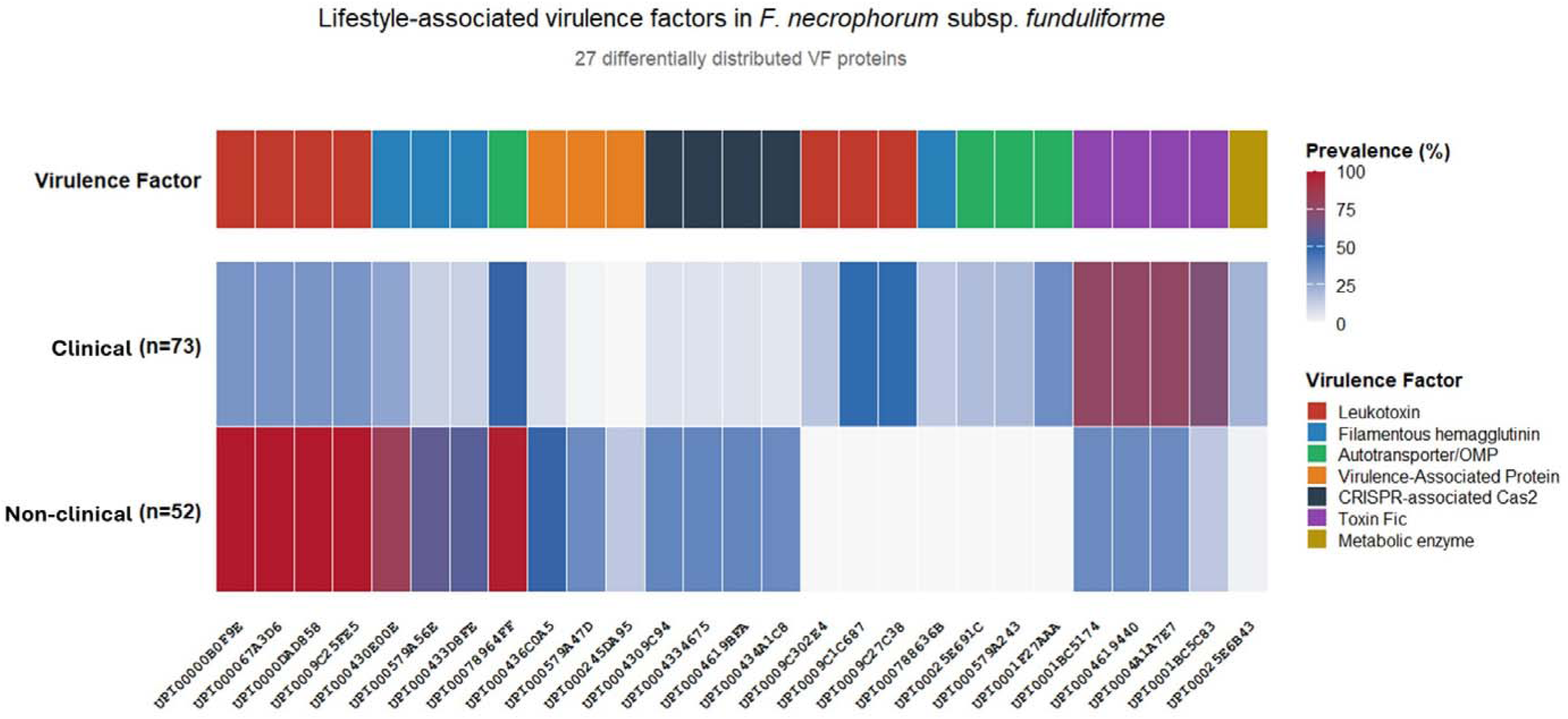
Lifestyle-associated virulence factor distribution in FNF strains. Heatmap visualization of 27 virulence factors (VF) showing significant differential distribution between clinical (diseased, n=73) and non-clinical (healthy, n=52) FNF strains. The top annotation bar indicates the virulence factor functional category for each VF (UniParc ID). The heatmap displays prevalence percentages using a blue-to-red diverging scale (0–100%), where red indicates higher prevalence and blue indicates lower prevalence. Non-clinical-associated factors (n=15) show higher prevalence in healthy strains, while pathogenic-associated factors (n=12) show higher prevalence in diseased strains. All comparisons achieved statistical significance (Fisher’s exact test with FDR correction, q < 0.05). Color scale represents prevalence (%): white denotes 0%, blue denotes intermediate (∼50%), and red denotes 100%, with color intensity proportional to prevalence values.

**Figure 7:**
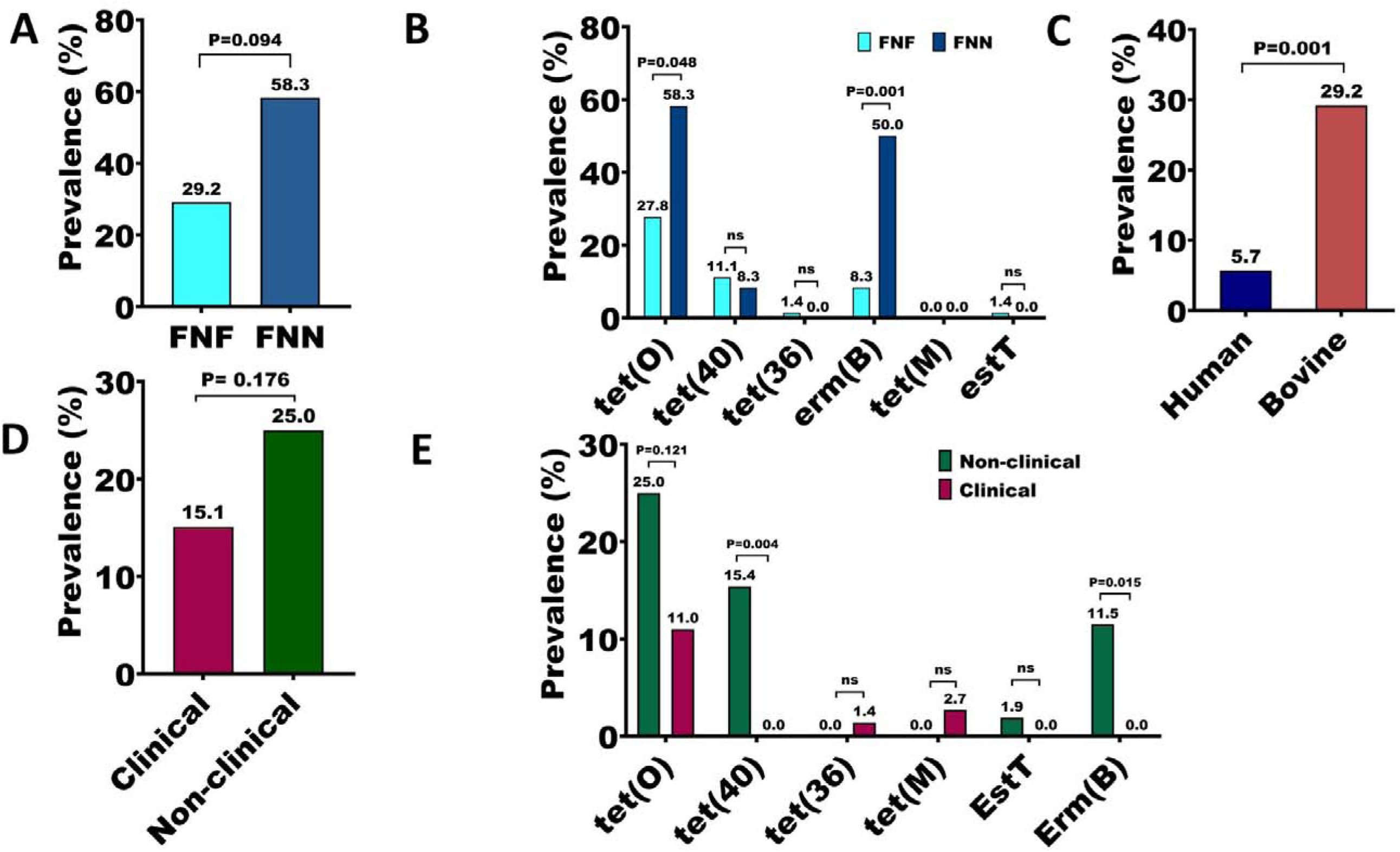
Antimicrobial resistance gene (ARG) prevalence across *Fusobacterium necrophorum* subspecies, host species and pathogenicity status. (A) Overall ARG prevalence (%) in *F. necrophorum* subspecies, comparing *F. necrophorum* subsp. *funduliforme* (FNF; n = 125) and *F. necrophorum* subsp. *necrophorum* (FNN; n = 12). **(B)** Prevalence of individual ARGs by subspecies. **(C)** ARG prevalence (%) by host species (bovine n = 84, human n = 53). **(D)** Comparison of ARG prevalence (%) between non-clinical (n = 52) and pathogenic (n = 20) bovine FNF strains. **(E)** Gene-specific prevalence (%) in non-clinical versus pathogenic bovine FNF strains. Statistical significance was assessed using Fisher’s exact test; **ns** indicates not significant.

**Table 3:**
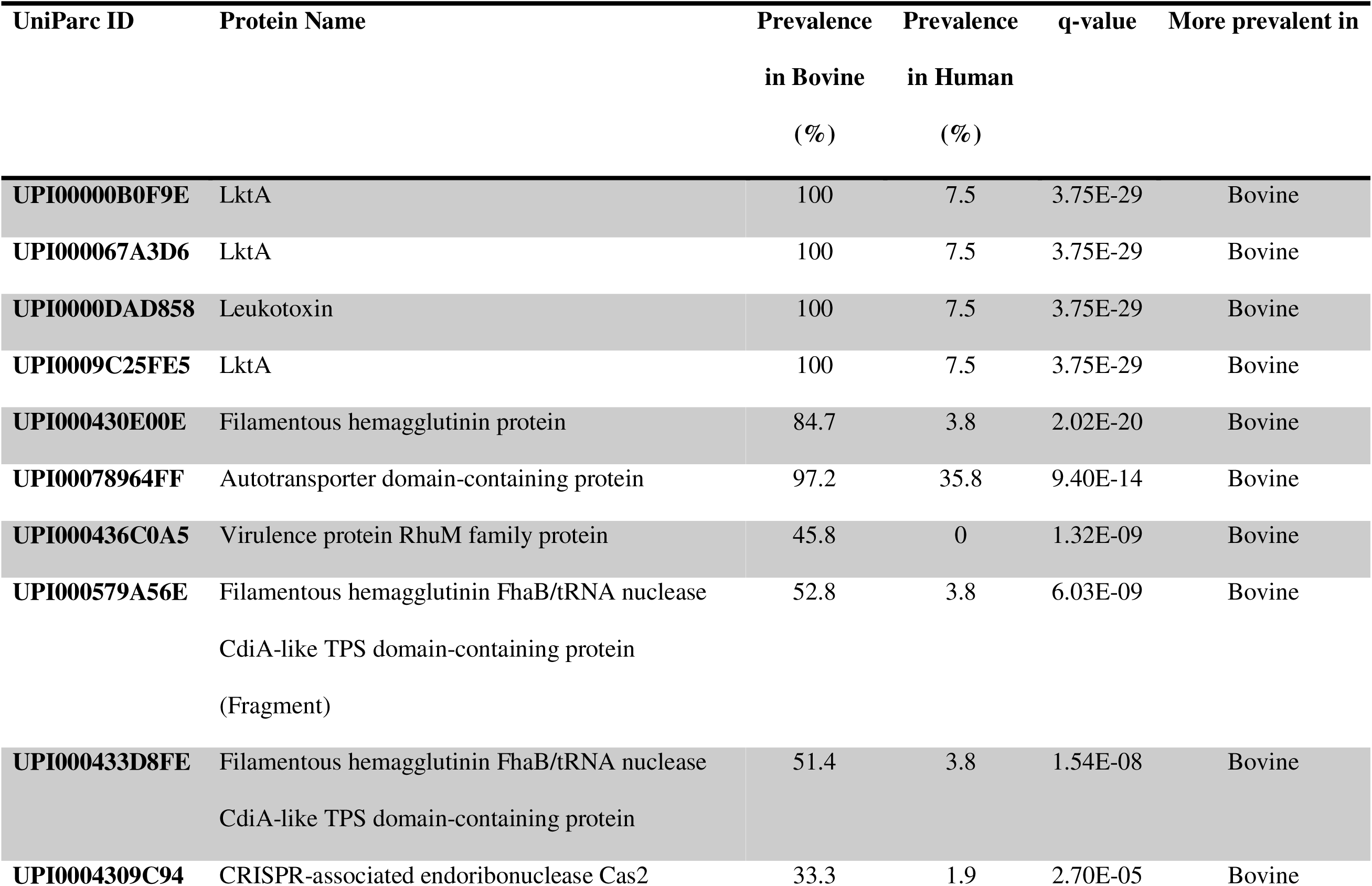
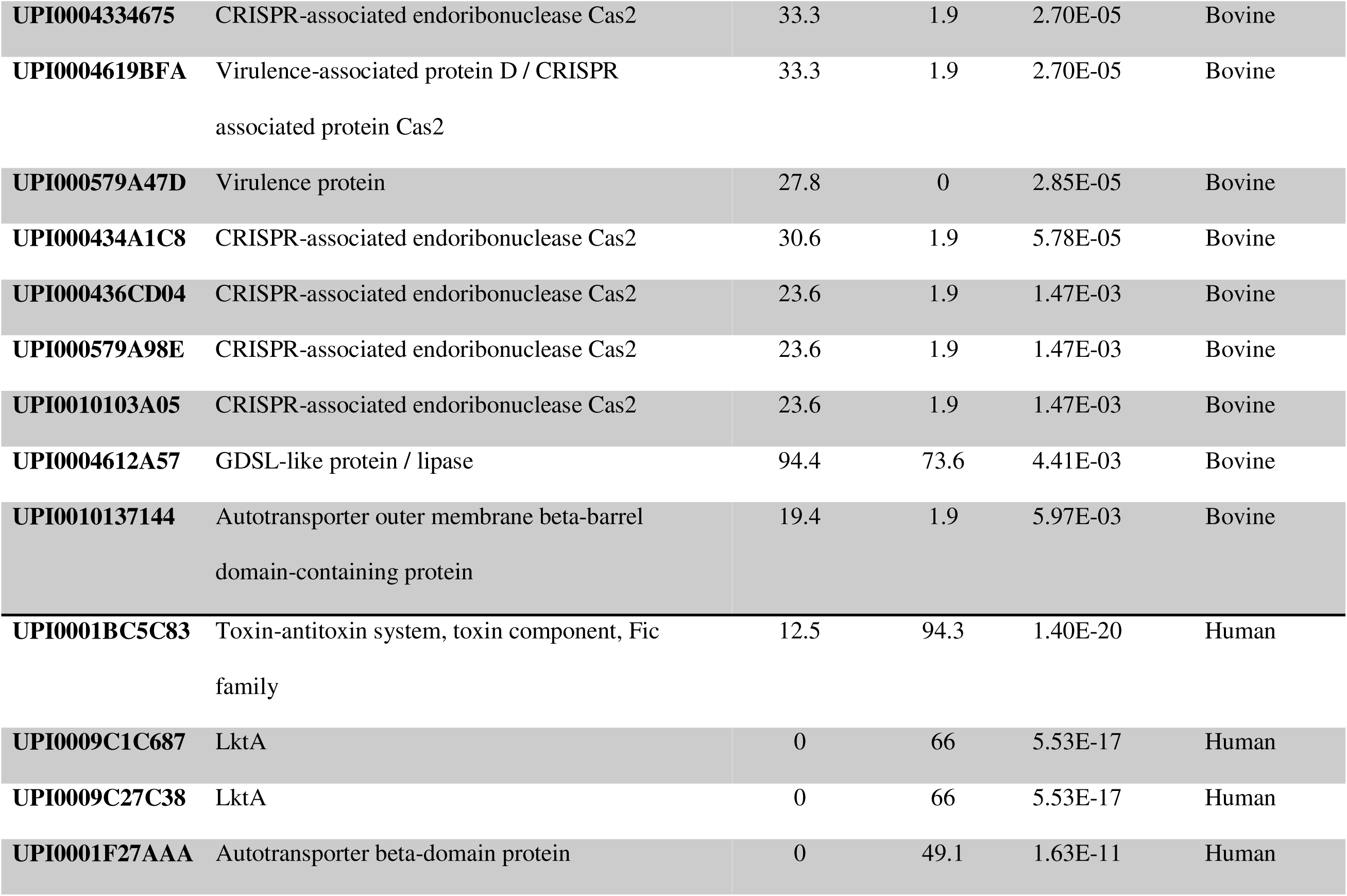
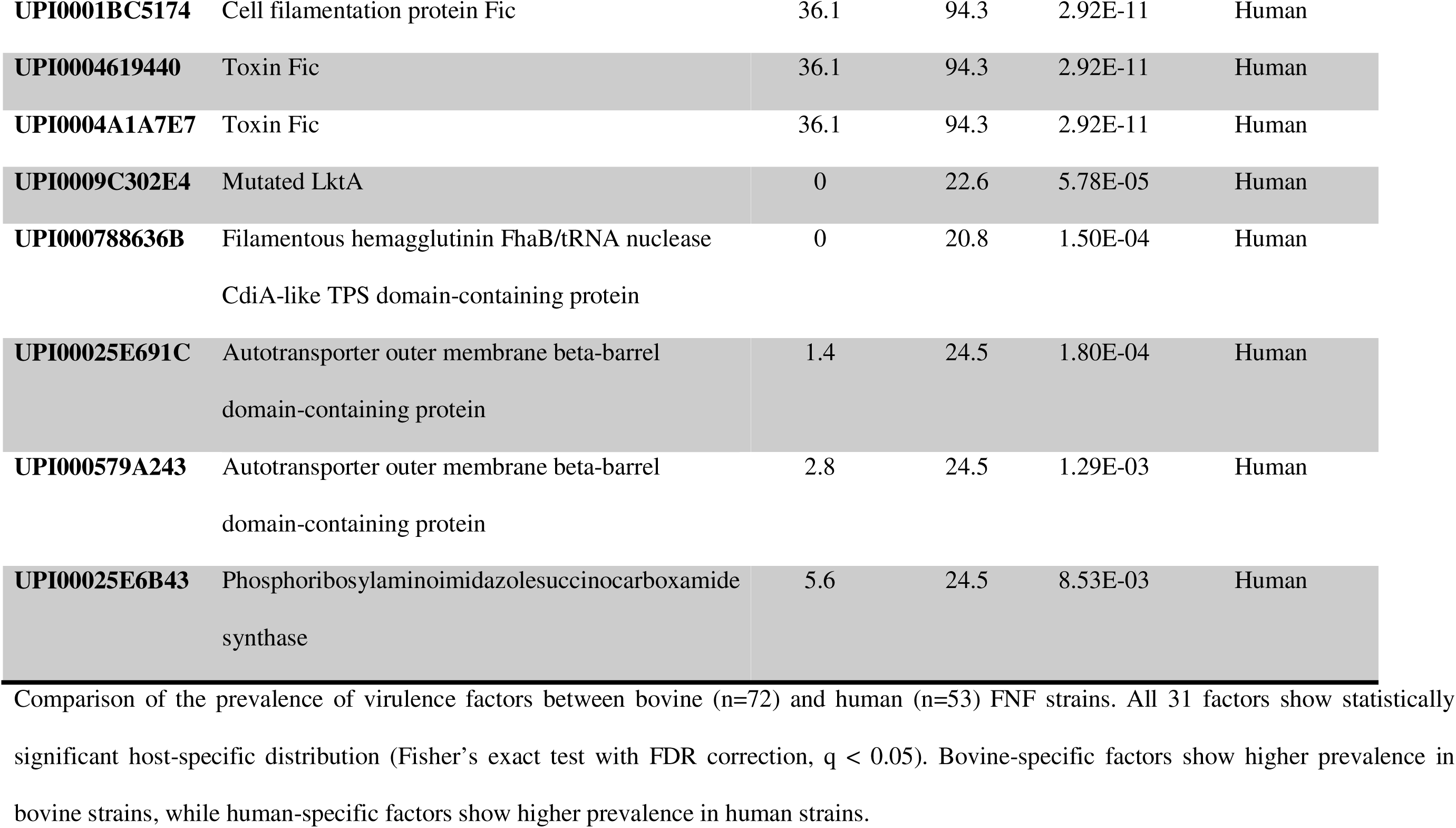
Host-associated distribution of virulence factors in *Fusobacterium necrophorum* subsp. *funduliforme* (FNF) strains.

Across both the bovine and human FNF strains, non-clinical strains showed higher ARG carriage than clinical strains (25.0% vs 15.1%) (Fig. 6E). The *tet*(O) gene was significantly more prevalent in non-clinical strains (25.0% vs 15.4%, *P* = 0.004) than clinical strains, and *estT* was detected only in non-clinical strains (11.5% vs 0%, *P* = 0.015) (Fig. 6E). Similar patterns were observed in bovine strains. Host species were also associated with ARG distribution, with bovine strains showing significantly higher ARG prevalence compared to human strains (29.2% vs 5.7%, *P* = 0.001) (Fig. 6C). This pattern was primarily driven by *tet*(O), which was significantly more prevalent in bovine strains (35.0% vs 3.8%, *P* = 0.002) (Fig. 6D). Among human strains, ARGs were largely confined to tonsillitis cases.

Among all the *F*. *necrophorum* strains (n = 137), comparison by anatomical origin showed that mediastinal lymph nodes (100%, 5/5), laryngeal (calf diphtheria; 100% *tet*(O) carriage, 7/7), and foot (foot rot;100%, 2/2) had the highest ARG prevalence, followed by bovine lung strains (66.7%, 6/9) (Table 3). None of the 25 seminal strains carried ARGs. Several strains encoded multiple ARGs, with 19 genomes (13.8%) harboring more than 2 ARGs on the same contig (Supplementary Fig. S4). For example, mediastinal lymph node isolate FNF104 harbored four resistance determinants [*tet*(O), *tet*(40), *erm*(B) and *estT*]. Isolate FNF103 carried a three-ARG combination [*tet*(O), *tet*(40) and *erm*(B)], while strain FNN107 from bovine foot rot tissue harbored *tet*(O) and *tet*(40). Bovine lung strains FNF91, FNF92, and FNF93 exhibited identical ARG profiles consisting of *tet*(O), *tet*(40), and *erm*(B) (Table 4).

**Table 4:**
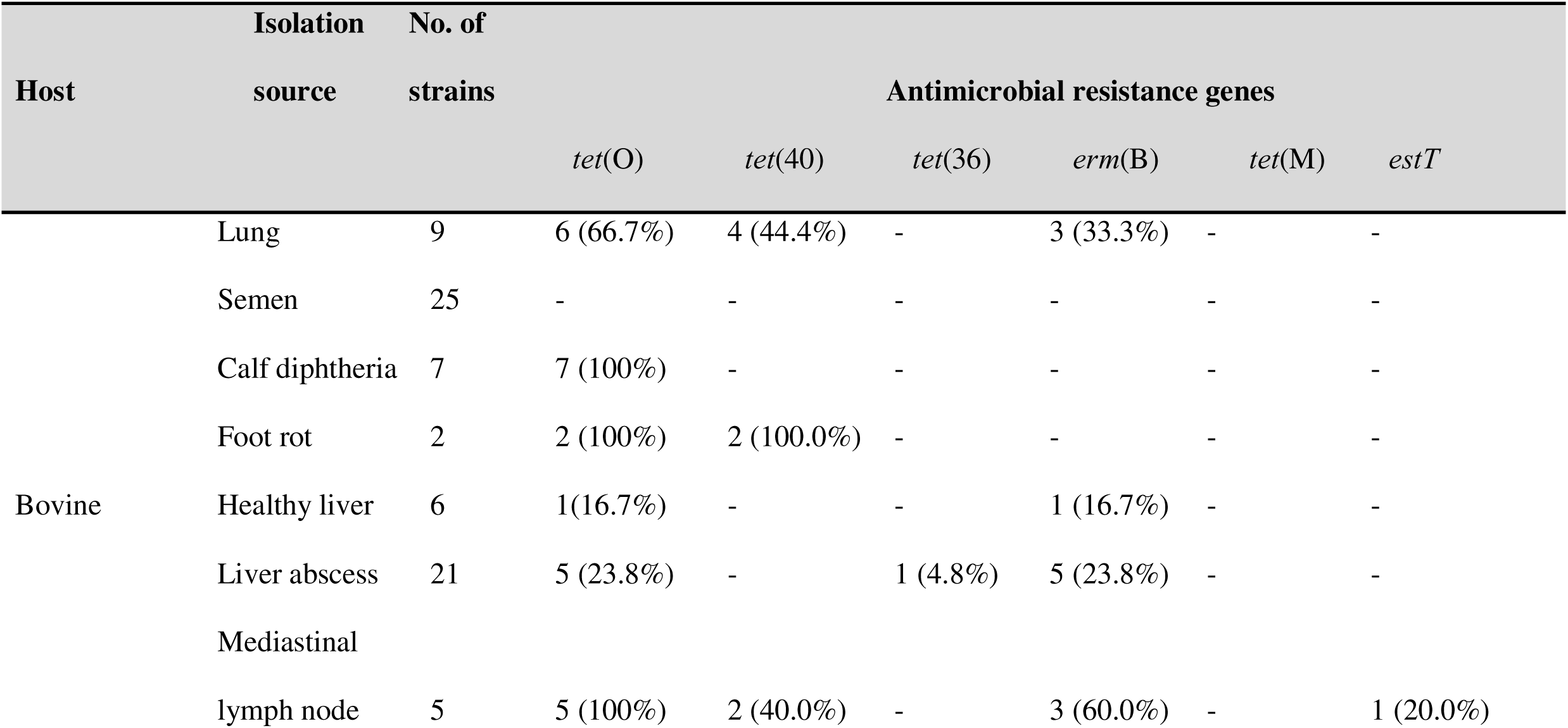
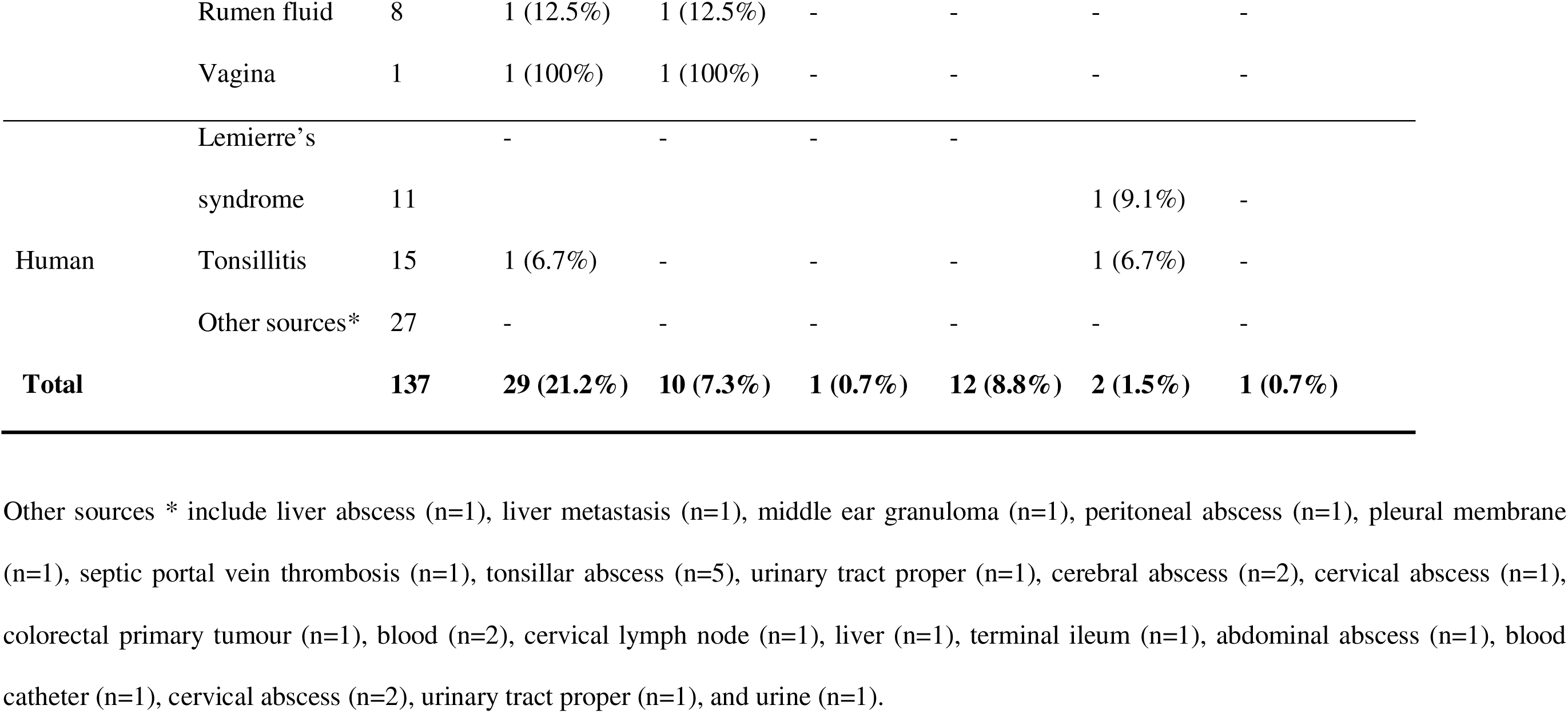
Prevalence (%) of antimicrobial resistance genes in *Fusobacterium necrophorum* (n = 137) by host and isolation source.

The MOB-suite analysis detected plasmid-associated sequences in 21 *F. necrophorum* genomes (15.3%), comprising predominantly chromosomally integrated mating pair formation type T (MPF_T) systems (18 genomes) and mobilization protein family V (MOBV) elements (3 genomes). Of Among these, only the human pharyngeal isolate AJ79 (*F. necrophorum* subsp. *funduliforme*) from a tonsillitis case harbored a confirmed circular extrachromosomal plasmid, a 9,887 bp element (pAJ01) that lacked ARG.

## DISCUSSION

To the best of our knowledge, this is the first study to utilize comparative genomics to determine whether subspecies identity, *necrophorum* vs. *funduliforme*, host species (bovine vs. human), clinical and non-clinical and anatomical sites (gastrointestinal vs. reproductive vs. respiratory tracts) are associated with genomic signatures that could help explain persistence of *F. necrophorum* as both a non-clinical commensal and an opportunistic pathogen. We analyzed 137 genomes (80 newly sequenced and 57 publicly available), including FNF (n = 125) and FNN (n = 12) strains from human clinical cases and bovine hosts. These *F. necrophorum* strains were originated from seven anatomical niches, including semen, liver (healthy and abscessed), lungs, ruminal fluid, larynx, hoof, and mediastinal lymph nodes. Overall, we observed that FN is genomically structured by subspecies, while ecological characteristics are further shaped by host- and niche-associated variation in accessory gene content, functional capacity, virulence architecture, and antimicrobial resistance.

Phylogenetic analysis of 137 FN genomes resolved two well-defined clades corresponding to the FNN and FNF subspecies with FN. Intra-subspecies comparisons revealed high genomic similarity within each group, with slightly higher ANI among FNN strains (99.31% ± 0.05%) than among FNF strains (98.63% ± 0.01%), suggesting that the FNN strains may be more genetically homogeneous than FNF (Bista et al., 2022). This genomic separation is consistent with earlier comparative genomic and ribotyping studies that reported subspecies-specific gene repertoires and evolutionary divergence between FNN and FNF strains (Okwumabua et al., 1996; Narongwanichgarn et al., 2003; Bista et al., 2022).

Phenotypic differences between FNF and FNN subspecies have also been described previously, including differences in leukotoxin production and virulence potential (Tadepalli et al., 2008b; Pillai et al., 2021). The higher intra-subspecies conservation observed for FNN strains relative to that of FNF strains suggests that FNN genomes may represent a more genetically conserved lineage, whereas genomes of FNF strains appear to encompass greater genomic diversity. In the present study, this pattern aligned with clear ecological differences between the subspecies. The FNF strains displayed a broader host and niche distribution, occurring in both bovine and human hosts across different anatomical sites including clinical and non-clinical niches. In contrast, FNN strains were recovered exclusively from bovine hosts and were predominantly associated with disease, especially liver abscesses. Thus, while these findings suggest that FNF may have diverse accessory genomic adaptations enabling occupation of broader ecological niches, whereas FNN appears more closely associated with bovine hosts, expanded sampling of FNN strains is needed to confirm these subspecies-specific patterns.

Importantly, neither disease status nor anatomical source clearly structured the phylogeny of FN strains. Clinical and non-clinical *F. necrophorum* strains were distributed in both subspecies clades, and human clinical presentations such as tonsillitis, Lemierre’s syndrome, and abscesses did not result in distinct phylogenetic clusters. This indicates that pathogenicity in *F*. *necrophorum* may not be explained solely by either subspecies identification or colonization site. Instead, disease appears to arise from interactions among genomic characteristics, host environment, ecological niche, and dysbiosis within the resident microbial community (Nagaraja et al., 2005; Tadepalli et al., 2009; Carrara et al., 2024). This interpretation is consistent with recent whole-genome sequencing analyses of *F. necrophorum* strains associated with human *F. necrophorum* infections, which have shown that virulence-associated gene profiles tend to follow phylogenetic structure rather than clinical syndrome classification (Carrara et al., 2024).

Based on the 137 genomes analyzed in the present study, the *F. necrophorum* pangenome was open, consisting of 5,820 predicted genes. Open pangenomes are characteristic of bacterial species undergoing repeated ecological transitions and retaining adaptive flexibility through gene gain and loss mediated by recombination and horizontal gene transfer (HGT) (Mira et al., 2010; Francino, 2012; McInerney et al., 2017). Such genomic plasticity has been reported to enable rapid adaptation to changing hosts and environmental conditions while maintaining a conserved core genome (Carrara et al., 2024). Likewise, previous comparative genomic analyses of FN strains (n=167) have reported that recombination within the accessory genome contributes substantially to strain divergence and subspecies differentiation (Bista et al., 2022). At the broader taxonomic scale, recent genus-wide genomic studies of *Fusobacterium* have revealed diverse genome architectures shaped by recombination and gene flow, indicating the potential role of HGT in the evolution of *Fusobacterium* genomes (Stott and Bobay, 2020; Bista et al., 2022; Molteni et al., 2024). The open pangenome observed in our present study therefore provides a genomic basis for the ecological versatility of *F. necrophorum*, which can colonize multiple host species and a wide range of anatomical environments.

Functional genomic profiling of the 137 FN genomes revealed substantial metabolic divergence between subspecies regardless of host origin, with 80 KOs differing significantly between FNN and FNF strains. The FNN genomes were dominated by carbohydrate transport and advanced glycation end-product degradation pathways, including raffinose/stachyose transporter components and fructoselysine metabolism. These functions may confer metabolic advantages in inflamed host tissues, where elevated glucose promotes protein glycation and generates fructoselysine and related AGEs that FNN can catabolize via the *frl* pathway (Francino, 2012; Kuczyńska-Wiśnik et al., 2025). This is consistent with the frequent recovery of FNN from lesions associated with the rumenitis–liver abscess complex (Scanlan and Hathcock, 1983; Nagaraja and Chengappa, 1998; Tadepalli et al., 2009; Amachawadi and Nagaraja, 2022). Meanwhile, FNF genomes were enriched for threonate metabolism, hemolysin-associated systems, restriction–modification, and alcohol dehydrogenase functions. These genomic features observed in FNF strains suggest distinct metabolic and membrane-interaction mechanisms that may influence FNF’s ecological distribution and tissue tropism (Zhang et al., 2006; Riordan, 2007; Bista et al., 2022; Molteni et al., 2024).

Comparison of non-clinical and clinical FNF strains revealed 60 significantly different KOs. Non-clinical strains showed higher abundance of cytochrome bd oxidase components, CRISPR-Cas systems, DNA methylation, and type III restriction enzymes, consistent with adaptation for persistence in complex microbial communities as compared to clinical FNF strains (Francino, 2012; Umaña et al., 2022; Beavogui et al., 2024). The clinical FNF strains displayed a higher prevalence of glutamate metabolism and specialized transport systems, potentially reflecting adaptation to nutrient conditions in damaged or inflamed host tissues as suggested previously (Nagaraja et al., 2005).

Within clinical FNF strains, 52 KOs differed significantly between bovine and human hosts. Bovine strains showed higher abundance of glutamate dehydrogenase, cytochrome bd oxidase, carbohydrate catabolic pathways, and CRISPR-associated systems. These results suggest adaptation to the polysaccharide-rich rumen and microaerobic conditions of liver abscesses (Gruninger et al., 2025). Human strains were characterized by higher abundance of NAD biosynthesis, nucleoside transport, DNA replication initiation, and osmolyte transport pathways, suggesting FNF’s adaptation strategy to nutrient-limited conditions of the pharynx and bloodstream (Groth et al., 2021; Bista et al., 2022; Carrara et al., 2024). These results suggest that although FNF colonizes both bovine and human hosts, it may rely on distinct functional repertoires in each ecological context. This supports the notion that bacterial phenotypes may arise from the interaction between lineage background and host-specific selective pressures (Read and Massey, 2014; Carrara et al., 2024).

The FNF strains isolated from the semen exhibited high genomic similarity based on ANI, phylogenetic clustering, and functional gene content. This phylogenetic trajectory suggests that the bovine reproductive tract may act as a selective ecological filter, shaping a distinct and recurrent *F. necrophorum* population adapted to this niche (Carrara et al., 2024; Kilama et al., 2025). To the best of our knowledge, this is the first study to perform whole-genome sequencing of *F. necrophorum* strains from healthy bull semen. However, this homogeneity may reflect the limited geographic origin of the strains, and broader sampling is needed to confirm this genomic pattern in semen-associated FNF strains (Kilama et al., 2025). These seminal FNF strains were enriched for genome maintenance functions, including *parB* and *ppa,* and lacked detectable ARGs, indicating that persistence in this niche may depend more on ecological compatibility than on resistance or virulence traits (Tadepalli et al., 2009; Carrara et al., 2024). This observation is consistent with our previous reproductive microbiome studies, in which we reported *F. necrophorum* in healthy bull semen, and with findings linking *Fusobacterium* abundance in bovine female reproductive tract to pregnancy outcomes (Webb et al., 2023a; Webb et al., 2023b; Kilama et al., 2025). Accordingly, we speculate that *Fusobacterium* may have a potential role as commensal in modulating the uterine microenvironment to support reproductive success. Thus, the present findings based on whole genome sequencing and comparative genomics further support the hypothesis that some FNF strains present in the male reproductive tract may function as commensals, with possible implications for both make and female fertility outcomes.

The virulence factor analysis further supports the view that pathogenic potential in *F. necrophorum* is modular rather than binary. We identified 84 distinct virulence factors distributed across all 137 FN genomes analysed. These include autotransporters, outer membrane proteins, hemagglutinins, leukotoxins, LPS/LOS biosynthesis enzymes, CRISPR-Cas-associated systems, and toxin-associated proteins. This virulence repertoire aligns with previous studies of *F. necrophorum* pathogenesis emphasizing the combined roles of leukotoxin activity, adhesion, invasion, and immune evasion rather than a single dominant determinant (Tan et al., 1996; Nagaraja et al., 2005; Ludlam et al., 2009; Carrara et al., 2024). However, only about one-third of these variants were part of the core repertoire, while the remainder were accessory or rare, indicating that much of the virulence-associated repertoire in this species is lineage dependent.

The differences between the genomes of FNN and FNF strains were particularly informative. Among the bovine derived strains, FNN was enriched for invasion-associated factors including *fadA* adhesin, a GDSL lipase/esterase, and a serine protease, meanwhile FNF was enriched for autotransporter outer membrane proteins. At the same time, several virulence genes, including leukotoxin components and LPS/LOS-associated genes, were conserved across both subspecies, suggesting a shared set of core pathogenicity-associated features alongside subspecies-specific determinants that may influence host interaction and tissue invasion (Carrara et al., 2024). Previous studies showed that adhesion of *F. necrophorum* to bovine endothelial cells is mediated by outer membrane proteins and is stronger in FNN than in FNF, with additional adhesion-associated proteins identified in recent genomic studies (Kumar et al., 2013b; Kumar et al., 2015; He et al., 2022; Bista et al., 2023).

When we compared non-clinical with clinical FNF strains, the non-clinical mainly exhibited higher abundance for several leukotoxin variants, whereas clinical FNF strains showed enrichment for Fic domain-containing toxin-antitoxin (TA) systems. Fic toxins catalyze AMPylation of host Rho GTPases, impairing GTP-dependent cytoskeletal signalling and innate immune activation (Roy and Cherfils, 2015; Veyron et al., 2018). The association of Fic TA systems with clinical niche specialization has been documented in other bacterial pathogens such as *Escherichia coli*, where they contribute to host tissue persistence and immune evasion (Harms et al., 2016; Sonika et al., 2023). Thus, the enrichment of Fic toxins in human FNF strains may reflect human host-adapted virulence strategies underlying invasive diseases and clinical presentations.

The presence of leukotoxin variants in non-clinical FNF strains highlights that pathogenic potential in *F. necrophorum* may likely depend not only on virulence genes presence but also on allelic variation and genomic regulations (Holm et al., 2017; Carrara et al., 2024). Indeed, multiple leukotoxin variants have been described in human-derived FNF strains (Holm et al., 2017). The specific virulence-associated profiles have recently been linked to invasive disease like Lemierre’s syndrome (Holm et al., 2017; Carrara et al., 2024). The host-specific virulence patterns within FNF strains further indicate the existence of adaptive diversification within the FN species. Bovine FNF strains showed higher enrichment of leukotoxin variants, filamentous hemagglutinins, and autotransporter proteins, whereas human strains showed a higher prevalence of Fic toxin–antitoxin systems and alternative leukotoxin variants. Accordingly, it is evident to speculate that FNF may comprise host-associated subpopulations that rely on different virulence strategies, with adhesion- and membrane-associated factors appearing more prominent in bovine strains and toxin-associated systems more common in human strains (Tadepalli et al., 2008a; Holm et al., 2017; Carrara et al., 2024). In addition, several virulence factors associated with autotransporter proteins, toxins, hemagglutinins, and LOS biosynthesis enzymes showed strong anatomical-site specificity within the FNF strains. We also observed that the autotransporter proteins were enriched in FNF strains isolated from the upper respiratory and thoracic tract (calf diphtheria, mediastinal lymph nodes, and lungs) but absent in strains derived from semen and ruminal fluid. In contrast, filamentous hemagglutinins and LOS-associated genes were prevalent in seminal and healthy liver-origin strains. Therefore, our findings suggest that distinct selective pressures operate across bovine niches, including differences in epithelial contact, immune exposure, oxygen availability, and microbial competition. These pressures likely shape the ecology of FN, reinforcing the importance of adhesion and host–cell interactions in its niche adaptation and persistence(Kumar et al., 2013b; Kumar et al., 2015; Bista et al., 2023).

None of the FNF strains isolated from bull semen carried ARGs, consistent with the low prevalence across *F. necrophorum* and likely because the FNF strains originated from yearling bulls with minimal exposure to antibiotic treatment. In addition, seminal FNF strains exhibited reduced virulence factor profiles, lower mobile genetic elements, and enrichment of CRISPR-Cas systems relative to non-semen derived clinical FNF strains. Given the proposed semen-to-cow-calf transmission pathway (Kilama et al., 2024; Kilama et al., 2025), this route is unlikely to contribute to resistance dissemination and highlights that seminal FNF strains as potential candidates for probiotic development in bovine reproductive health.

The distribution of ARGs in *F. necrophorum* appears to be influenced by ecological context, particularly antimicrobial exposure in livestock production system (Nagaraja et al., 2005; Amachawadi and Nagaraja, 2022). The tylosin phosphate, a macrolide antibiotic, is the most commonly used in-feed antimicrobial in North American beef production for the control of liver abscesses caused by FN species. Whereas, tetracyclines have historically been used for liver abscess control, respiratory disease, and foot rot in feedlot cattle (Nagaraja and Chengappa, 1998; Nagaraja et al., 2005; Amachawadi et al., 2017; Davedow et al., 2020; Pinnell et al., 2022). The detection of *tet*(O), *tet*(40), and *erm*(B) predominantly in FN strains isolated from liver abscess and other clinical sources could be attributed to sustained antimicrobial selection pressure in feedlot systems, rather than representing a broadly distributed species-wide trait.

The co-location of *erm*(B), *tet*(40), and *tet*(O) with IS3 family transposase elements in FNF104 and FNF103 suggests the potential acquisition of resistance via mobile genetic elements. Horizontal gene transfer (HGT) within animal-associated bacterial population is well documented as a major driver of antimicrobial resistance emergence in livestock environments (Didelot and Maiden, 2010; Francino, 2012). However, the absence of flanking transposase signatures in FNF91 and FNN107 strains carrying ARGs may likely indicate possible chromosomal stabilization over time after HGT (Arkhipova et al., 2023; Yaikhan et al., 2024). Given that tylosin is used in approximately 71% of U.S. feedlot cattle, raising concerns about ongoing selection and potential dissemination of macrolide and co-selected tetracycline resistance determinants within *F. necrophorum* populations and the broader feedlot microbiome (Lechtenberg et al., 1998; Sabino et al., 2019; Perry et al., 2023; Elbehiry et al., 2025).

Although the FN strains from bovine and human were mainly clustered within host-associated groups, several FN strains from both hosts were nearly identical based on ANI, consistent with shared or closely related strains. While this does not demonstrate zoonotic transmission, it suggests that some FN strains may circulate across host boundaries or arise from shared reservoirs (Riordan, 2007). This observation is particularly relevant given ongoing uncertainty regarding the origins of human *F. necrophorum* infections, including Lemierre’s syndrome, which may originate from the host microbiome, environmental exposure, or person-to-person transmission (Riordan, 2007; Perry et al., 2023).

There are few limitations of the current study that warrant some considerations. First, the number FNN strains was underrepresented relative to FNF strains, with only 12 strains available, including a single pathogenic strain from a bovine liver abscess. This limited sample size precluded robust assessment of differences in virulence factor distribution between clinical and non-clinical FNN strains. This imbalance stemmed from the inherent difficulty of culturing FNN strains, which is less prevalent and more fastidious than FNF strains, as observed previously (Kilama et al., 2025). Second, publicly available human genomes were exclusively clinical strains, limiting the representativeness of cross-host comparisons and potentially inflating associations between specific virulence factor profiles and disease. Third, the use of short-read Illumina assemblies constrained the resolution of mobile genetic elements and structural genomic variation. Moreover, these data do not capture regulatory variation, gene expression dynamics, or allele-specific functional effects, which may be particularly important for surface-associated proteins and leukotoxin variants (Holm et al., 2017; Carrara et al., 2024). Despite these limitations, our current study presents the first application of whole-genome sequencing and comprehensive comparative genomics to FN strains from human and bovine hosts across diverse clinical and non-clinical sources.

Future research should expand genomic sampling across diverse hosts, anatomical niches, and geographic regions, and incorporate long-read sequencing to resolve plasmids, mobile elements, and structural variation. Transcriptomic and functional studies will be essential to determine the regulation of key loci, including *lktA,* adhesion-related proteins, and metabolic pathways during colonization and disease. Culturomics, metabolic profiling, and molecular labeling approaches will also be important to track *F. necrophorum* transmission across anatomical sites, including potential cross-seeding from the rumen to the reproductive tract. Furthermore, longitudinal studies are needed to define its role in healthy reproductive microbiomes and effects on offspring immune development. Given their role in polymicrobial infections, future models incorporating microbial interactions will be critical to understanding clinical outcomes. Together, these priorities highlight the need for integrated One Health genomic surveillance to clarify cross-host transmission and shared reservoirs.

## Conclusion

Comparative genomic analysis of 137 *F. necrophorum* genomes (80 generated in our present study), spanning nine bovine anatomical sites and including human strains from clinical and non-clinical sources, revealed clear subspecies-level structure with limited clustering by disease status or anatomical niche. The *F. necrophorum* species possesses an open pangenome with extensive accessory gene diversity, consistent with ecological flexibility across hosts and tissues. Subspecies-specific metabolic signatures included enrichment of carbohydrate transport and fructoselysine metabolism in FNN, and threonate metabolism and hemolysin-associated functions in FNF strains. Virulence factors such as leukotoxin, hemagglutinins, and outer membrane proteins were broadly conserved, whereas other determinants varied by subspecies and niche. The ARGs were infrequent, largely restricted to tetracycline and macrolide resistance, and absent in seminal strains. Overall, these findings suggest that pathogenic potential in *F. necrophorum* is determined by the interplay between an open pangenome, subspecies-specific metabolic and virulence repertoires, host-associated genomic adaptation, and niche specialization.

## ACKNOWLEDGMENTS

We acknowledge the Center for Computationally Assisted Science and Technology (CCAST) at North Dakota State University for providing the high-performance computing resources used in this study. We thank the NDSU Veterinary Diagnostic Laboratory for providing *Fusobacterium necrophorum* strains from bovine foot rot cases. We also thank Godson Aryee, Md Shafinul Islam, and Kell Helmuth from Amat Lab NDSU, as well as Leigh Ann George and Xiaorong Shi from Kansas State University, for their assistance.

## AUTHOR CONTRIBUTIONS

Justine Kilama (Conceptualization, Funding acquisition, Data curation, Formal analysis, Investigation, Methodology, Software, Validation, Visualization, Writing—original draft, Writing—review & editing), Devin B. Holman (Data curation, Formal analysis, Software, Validation, Visualization, Writing—review & editing), Mina Abbasi (Methodology, Validation, Writing—review & editing), Raghavendra G. Amachawadi (Methodology, Resources, Writing—review & editing), Carl R. Dahlen (Investigation, Methodology, Validation, Writing—review & editing), T.G. Nagaraja (Conceptualization, Resources, Supervision, Writing—review & editing), and Samat Amat (Conceptualization, Data curation, Formal analysis, Funding acquisition, Investigation, Methodology, Project administration, Resources, Supervision, Validation, Visualization, Writing—original draft, Writing—review & editing).

## DATA AVAILABILITY

Raw sequencing data have been deposited in the NCBI Sequence Read Archive under BioProject accession PRJNA1446192

---

**Supplementary Figure S1.**
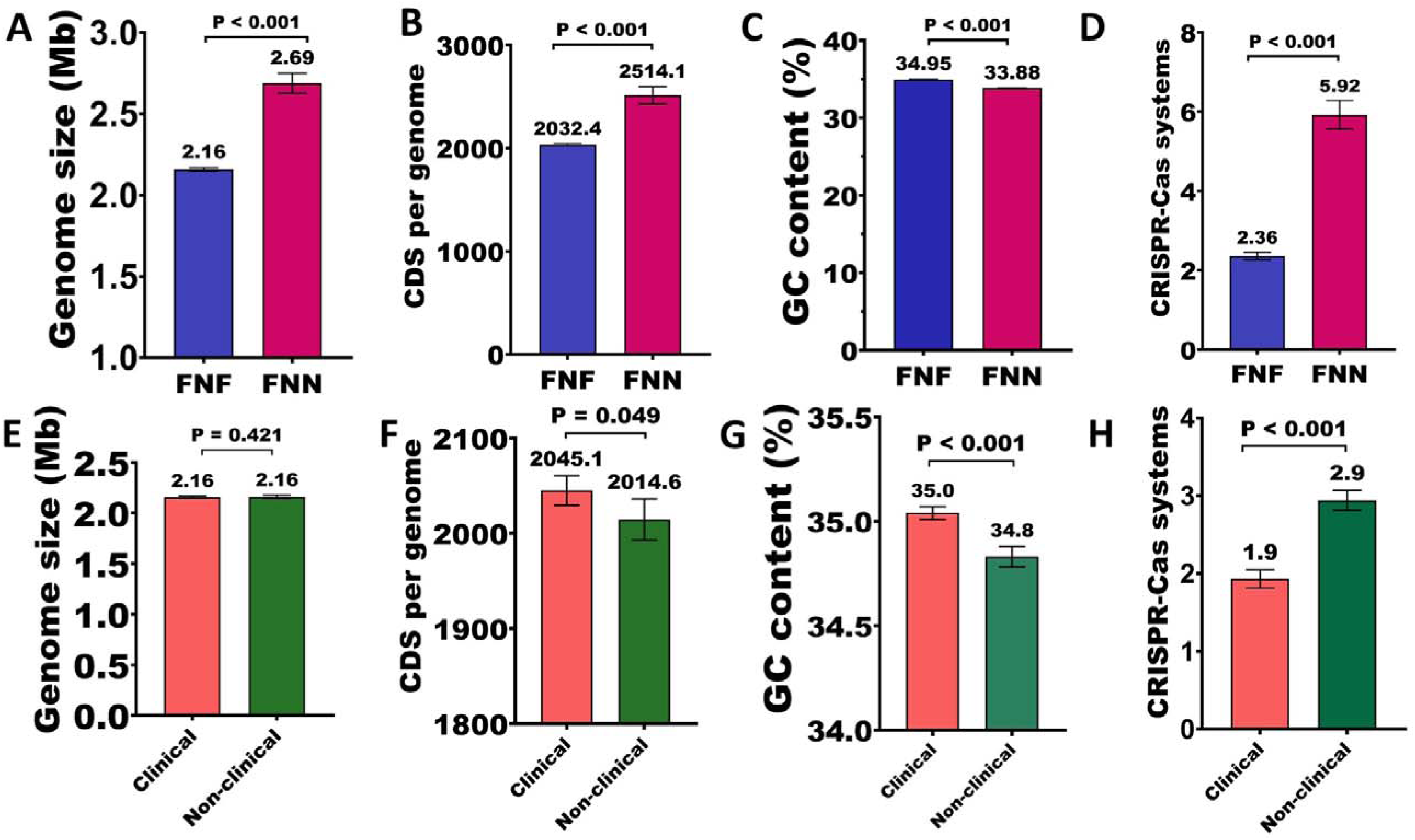
Subspecies and pathogenicity comparisons in *Fusobacterium necrophorum.* (A-D) Subspecies comparison between *Fusobacterium necrophorum* subsp. *funduliforme* (FNF) (n=125) and FNN (n=12) strains showing genome size (A), coding sequence count (B), G+C content (C), and CRISPR array abundance (D). (E-H) Pathogenicity comparison between pathogenic (n=73) and non-clinical (n=52) strains showing genome size (E), CDS count (F), G+C content (G), and CRISPR-Cas systems (H). Statistical significance determined by Mann-Whitney U tests. Error bars represent standard error of the mean.

**Supplementary Figure S2.**
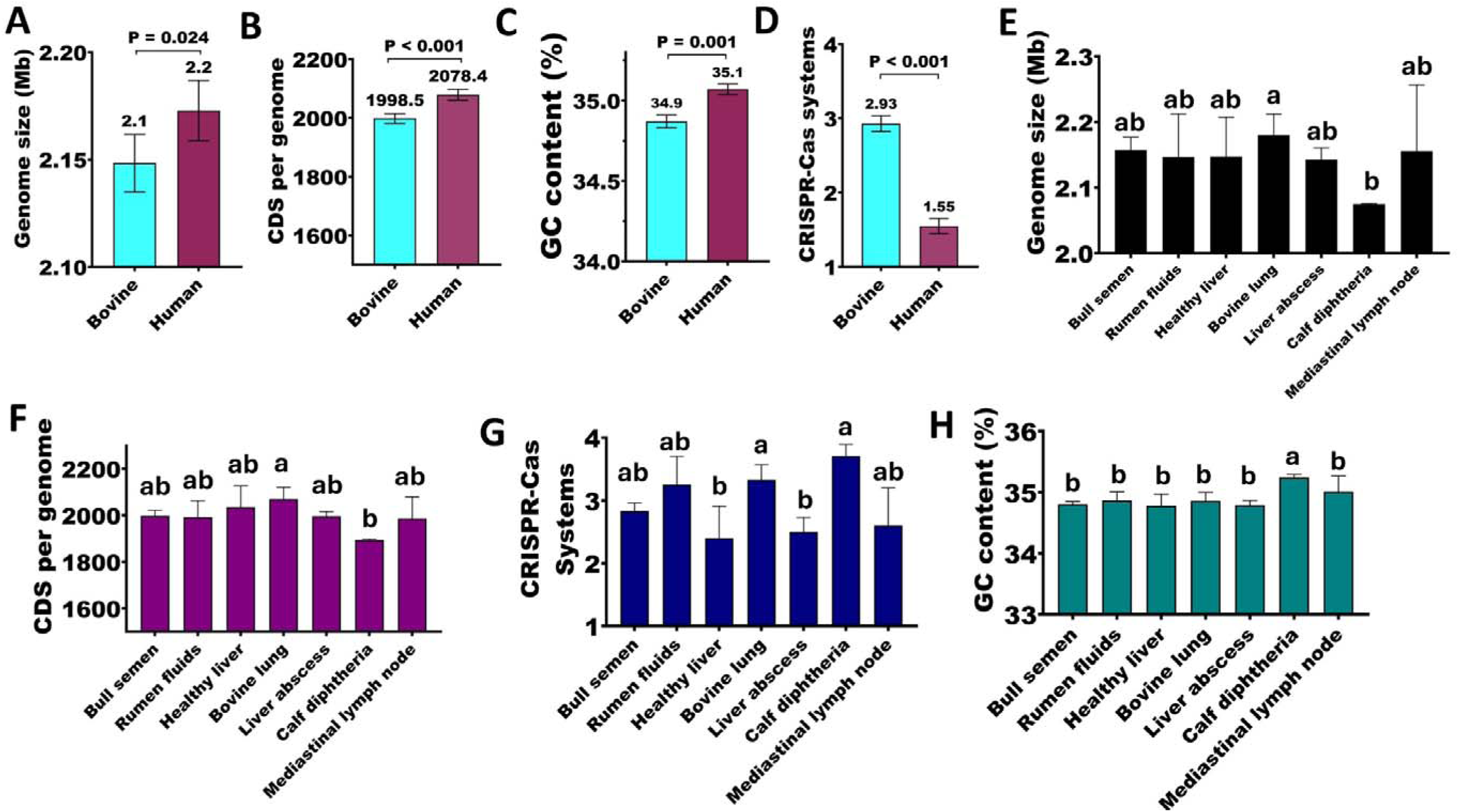
Host- and tissue-associated variation in genomic characteristics of *Fusobacterium necrophorum.* (A-D) Comparison of bovine (n=72) and human (n=53) strains Differences were assessed using Mann-Whitney U tests. **(E-H)** Comparison across anatomical isolation sources: bull semen (n=24), rumen fluid (n=8), healthy liver (n=5), bovine lung (n=9), liver abscess (n=12), calf diphtheria (n=7), and mediastinal lymph node (n=5), showing genome size (E), CDS count (F), CRISPR-Cas systems (G), and G+C content (H). Statistical significance was determined using Kruskal–Wallis tests followed by post hoc comparisons; groups not sharing letters (a, b) differ significantly (p < 0.05). Error bars represent the standard error of the mean.

**Supplementary figure S3.**
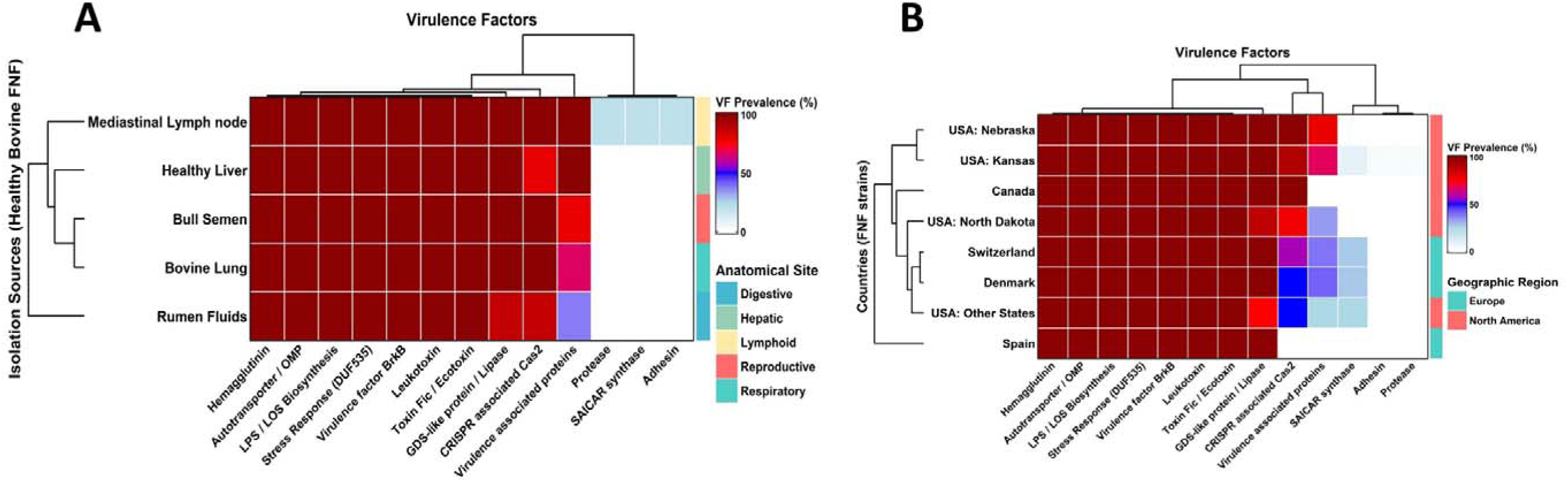
Virulence factor (VF) distribution in *Fusobacterium necrophorum* subsp. *funduliforme* (FNF) strains by anatomical site and geographic origin. (A) Heatmap showing the prevalence (%) of VFs in non-clinical bovine FNF strains across healthy anatomical sites (n=52). Rows represent isolation sources hierarchically clustered by similarity and columns represent VF categories. Color intensity indicates prevalence (red = high, blue = low). Anatomical sites are annotated by tissue type. (B) Heatmap showing the prevalence (%) of VFs across geographic origins (n=125). Rows represent countries clustered by similarity and columns represent VF categories. Countries are annotated by geographic region (Europe = teal, North America = red). Both analyses demonstrate variation in VF profiles associated with anatomical site and geographic regions.

**Supplementary figure S4:**
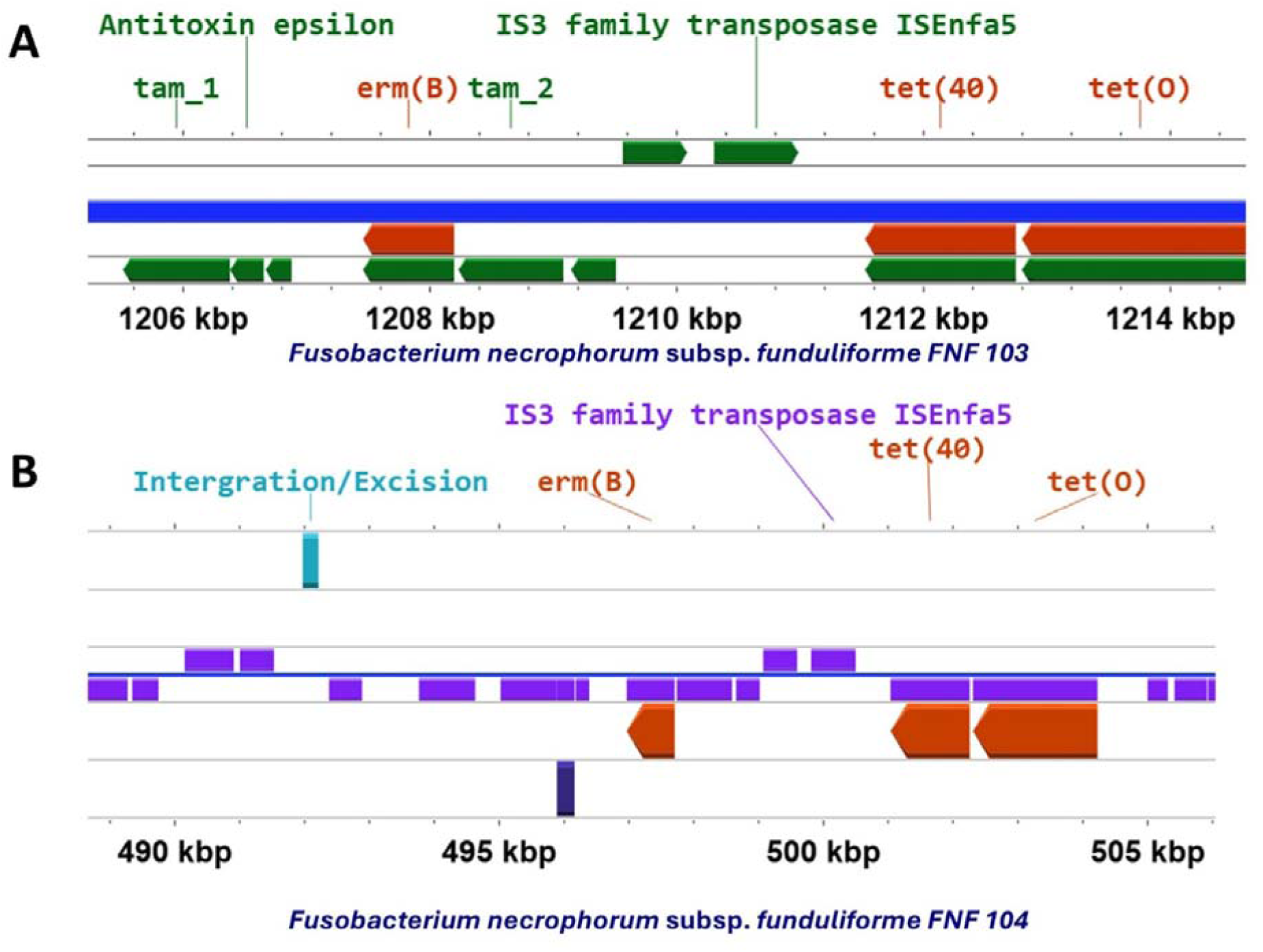
Chromosomal organization of antimicrobial resistance genes (ARGs) co-located on the same contig in *Fusobacterium necrophorum* genome assemblies. *F. necrophorum* subsp. *funduliforme*:(A) FNF103, (B) FNF104, Red arrows represent ARG sequences, green (A) and purple (B) arrows indicate other coding sequences as annotated by Prokka and visualized in ProkSee

